# A Shape Analysis Algorithm Quantifies Spatial Morphology and Context of 2D to 3D Cell Culture for Novel Quantitation of Phenotypes

**DOI:** 10.64898/2026.02.02.703425

**Authors:** David H. Nguyen, Michael Bruck, Jennifer M. Rosenbluth

## Abstract

**SUMMARY:** This article explains how novel morphological features in cells and organoids can be quantified using the linearized compressed polar coordinates (LCPC) transform, a spatial algorithm that captures properties that traditional metrics, such as area, volume, and surface area, cannot. Best practices for shape orientation and alignment are discussed.

Numerous studies have shown that the morphological phenotype of a cell or organoid correlates with its susceptibility to anti-cancer agents. However, traditional methods of measuring phenotype rely on spatial metrics such as area, volume, perimeter, and signal intensity, which work but are limited. These approaches cannot measure many crucial features of spatial context, such as chirality, a property of left- and right-handedness. Volume cannot register chirality because the left and right shoes hold the same volume. Though spatial context in the form of chirality, gravity direction, and polarity axis is intuitive to humans, the traditional metrics used by cell biologists, pathologists, radiologists, and machine learning practitioners to date cannot capture these fundamental notions. The Linearized Compressed Polar Coordinates (LCPC) transform is a novel algorithm that can capture spatial context unlike any other metric. The LCPC transform translates a two-dimensional (2D) contour into a discrete sinusoidal wave by overlaying a grid system that tracks the points of intersection between the contour and the grid lines. It turns the contour into a series of sequential pairs of discrete coordinates, with the independent coordinate (x-coordinate) being consecutive positions in 2D space. Each dependent coordinate (y-coordinate) consists of the distance between an intersection of the contour and gridline to the origin or baseline of the grid system. In the form of a discrete sinusoid wave, the Fast Fourier Transform is then applied to the data. In this way, the shapes of cells in 2D and 3D cell culture are systematically and multidimensionally represented, enabling robust quantitative stratification that will reveal insights into treatment resistance.

## INTRODUCTION

Three-dimensional (3D) organoid culture has proven superior to two-dimensional (2D) cell culture in mimicking *in vivo* biology^1,2^. Organoids have become indispensable in cancer research for screening for effective treatments and for gaining insights into disease progression^3–6^. While it is well known in cancer biology that distinctions in morphology correlate with distinct biological behavior of cells and tissues, the field relies on traditional shape metrics that are limited in scope. This study describes a novel spatial algorithm that objectively quantifies subtle morphological features with unprecedented precision compared to traditional approaches such as area, volume, and surface area.

The linearized compressed polar coordinates (LCPC) transform was invented to objectively and quantitatively describe complex morphologies observed in tissue histopathology, with the aim of enhancing the grading of colon polyps^7^. The desire to then expand this approach to capture macroscopic MRI-based spatial features in brain pathologies, such as bipolar disorder and Alzheimer’s disease, led to augmentations that enhanced it even further. This study describes the step-by-step protocol and best practices for applying two versions of LCPC transform: the parallel grid system and the radial grid system, both of which have open-source scripts and tutorial videos.

Aside from traditional geometric methods, there exist multiple abstractions that have been effective for measuring the complex shapes of cells and organs. These include measuring eccentricity, skewness, contrast, and kurtosis, along with Zernike moments for capturing circularity, asymmetry, edge irregularity, and global contour structure^8–10^. Fractal analysis has also been useful to reduce complex contours into scalar values, such as Fractal Dimension, which measures how irregular the contour is, and lacunarity, which measures how heterogeneously irregular gaps appear in the contour^11,12^. For measuring texture, the Gray Level Co-occurrence Matrix (GLCM) method is popular for measuring contrast, energy, homogeneity, correlation, and entropy^13,14^. While all of these methods are beneficial, none of them were designed to capture spatial context, such as the direction of gravity, left- vs. right-handedness, or the location of the central support structure that influences the direction of shape change. Furthermore, many of them produce one scalar value or just a handful of scalar values to represent spatial information, which is why the cited studies often use many of them in conjunction to assess complex shapes.

The LCPC transform was designed to produce many features from one measurement and to allow the addition of spatial markers that encode spatial context into the shape, such as the direction of gravity. The closest method to the LCPC transform^7^ was published a year afterwards^15^, sharing the same core idea as a starting point: apply the Fourier transform to cell contours to measure shape in the form of a frequency spectrum. However, the LCPC transform was independently developed to be applied via distinct grid systems that were meant to be used in conjunction with knowledge of the spatial context outside of the shape being measured. Furthermore, the inventor of the LCPC transform explains in this manuscript that the resulting frequency spectrum contains vast amounts of hidden spatial information. **Supplementary File 1** contains an extensive discussion of spatial contexts in biology that are often overlooked when using the cited methods and how the LCPC transform can be applied to capture this context. For readers whose shape data shows no difference between control vs. experimental groups, whether or not their eyes can see a difference in the shapes, or whose shape data shows very little difference, even though they suspect that there should be a bigger difference, they should try the LCPC transform.

The LCPC transform provides an unprecedented level of precision in measuring spatial information because it represents shapes in multiple dimensions. Unlike traditional methods, like area and volume, which yield only one scalar value per shape (i.e., 25 cm2), the results of the LCPC transform can yield multiple indices, each correlating with a different morphological aspect of a shape (i.e., roundness vs. sharpness of corners, smoothness vs. jaggedness of edges).

The LCPC transform describes a 2D shape by overlaying a grid of straight lines that intersect its contour. Each grid system has an origin (**Figure 1**) or a baseline (**Figure 2**) from which to measure linear distance. Refer to **Figures 1B** and **2B** for simplified flowcharts describing the algorithm. Every point of intersection between the gridlines and the contour of the shape is detected. The distance of each intersection is then calculated relative to a baseline or origin. In this way, the LCPC transform converts 2D shapes into a series of consecutive x and y pairs of coordinates. The x-coordinate represents the position of the gridline from zero to infinity, while the y-coordinate represents the distance of the intersection to the baseline or origin. In this form, which is a discrete sinusoid wave, the Fast Fourier Transform (FFT) is then applied to convert the data from the “position domain” to the frequency domain. If the x-coordinate represented time, then the “position domain” would be equivalent to the “time domain” in the standard applications of the FFT.

**Figure 1:**
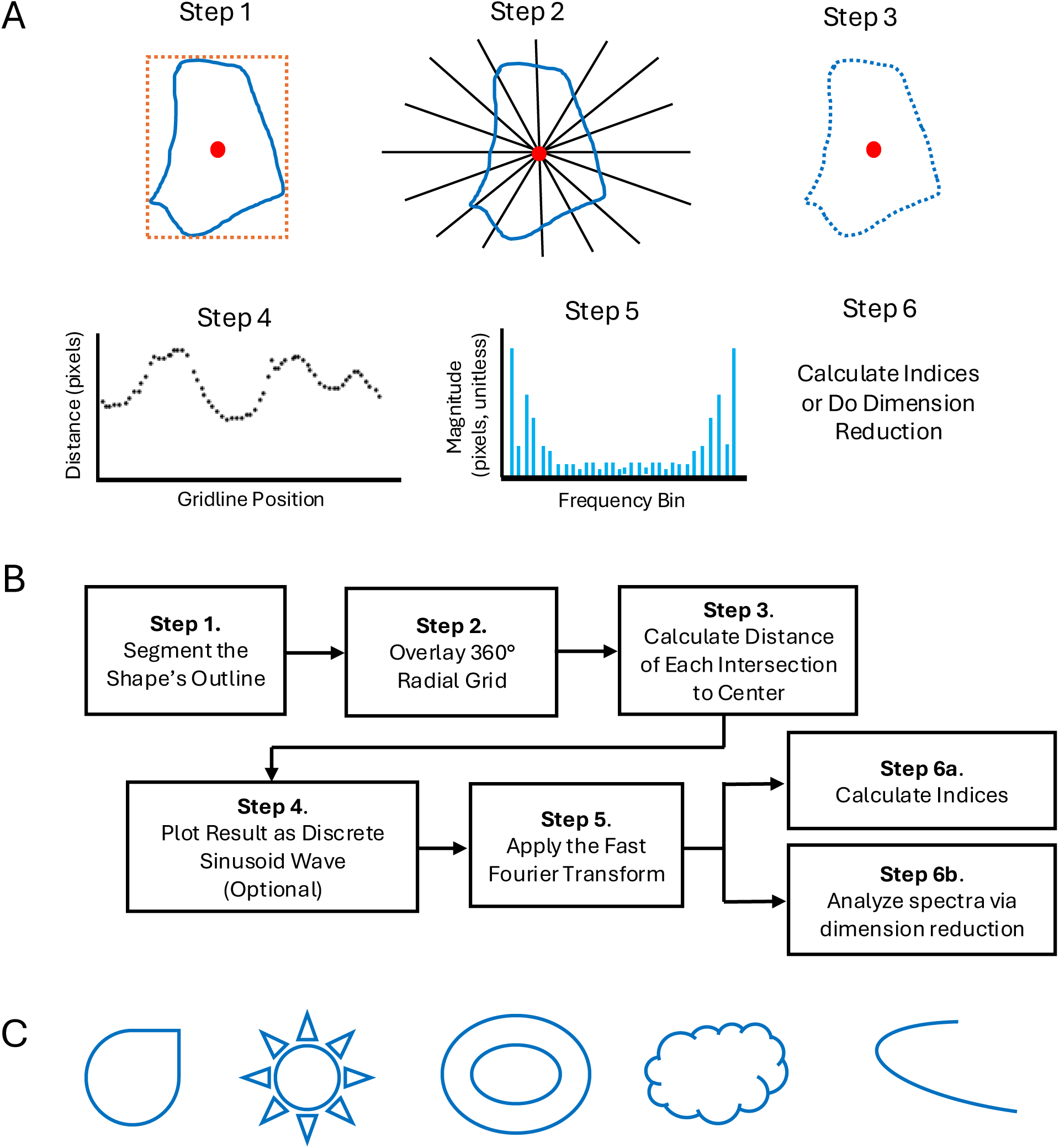
Description of the radial grid system of the LCPC transform. (**A**) Step 1: The shape of interest is segmented and extracted as a blue mask. Step 2: A bounding box (a.k.a. the smallest rectangle encasing and touching the object on all four sides) is determined for the mask. A system of radial grid lines originating from the center of the bounding box is overlaid onto the mask. The decision to pick the center of the bounding box instead of the centroid of the object as the origin was arbitrary. Step 3: The distance of each intersection between a gridline and the shape is calculated relative to the origin. Grid lines that have more than one intersection sum all those intersection distances into one value for that grid line. Step 4: The FFT is applied to turn the data in Step 3 into a frequency plot, which is a static multidimensional representation (Step 5) of the shape in Step 1. Step 6: A scalar value can be created from Step 5 via an index. See Figure 4 for a discussion on types of indices to calculate. (**B**) A flowchart showing a simplified description of the steps of the LCPC transform for the radial grid system. (**C**) Examples shape themes for which the radial grid system is optimal.

**Figure 2:**
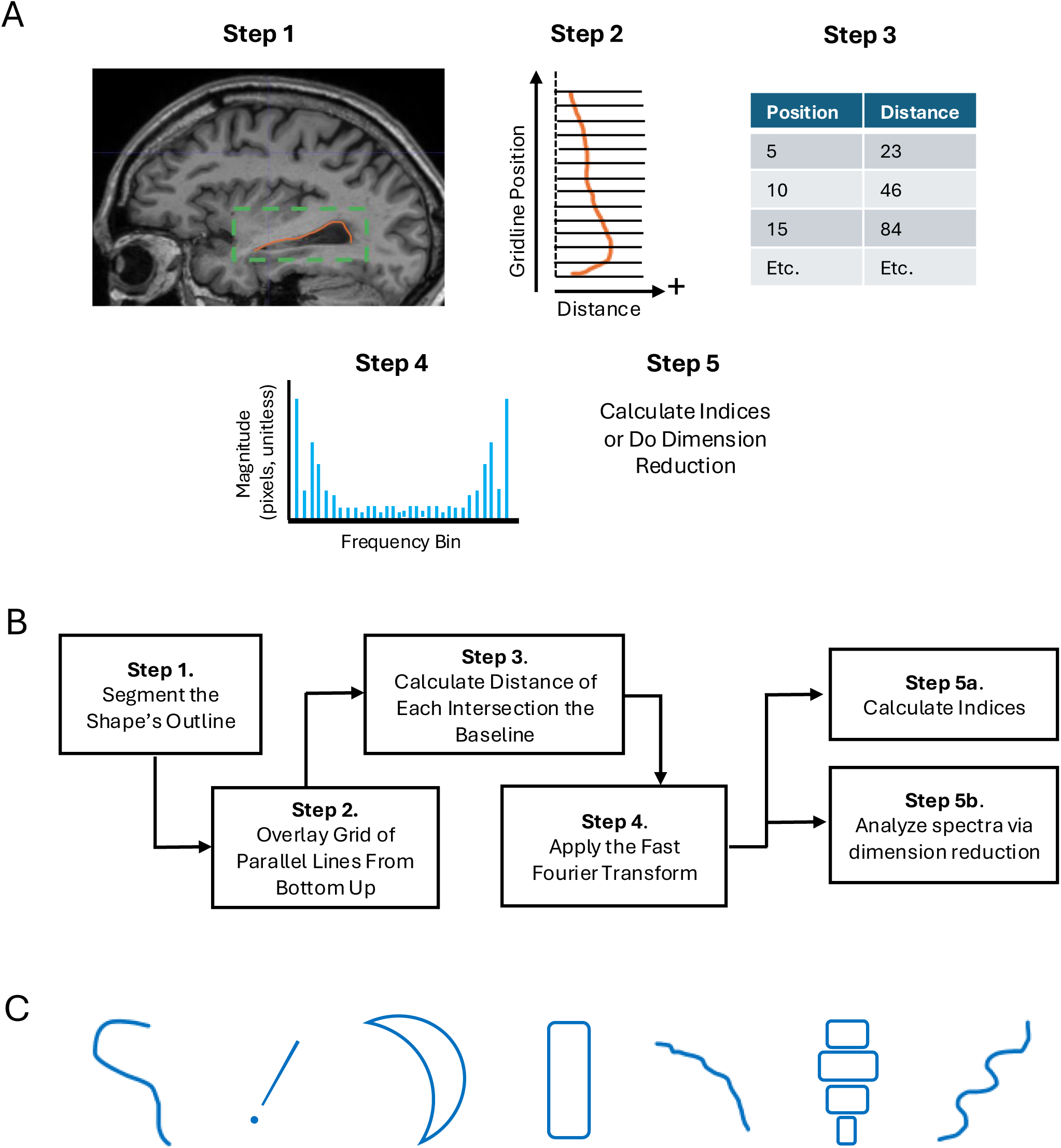
Description of the parallel grid system of the LCPC transform. (**A**) Step 1: The shape of interest is manually segmented and extracted as a binary mask. Step 2: A parallel grid system is overlaid onto the shape, which is rotated such that the base is horizontal. Step 3: The distance of each intersection between a gridline and the shape is calculated relative to the baseline. Grid lines that have more than one intersection sum all those intersection distances into one value for that grid line. Step 4: The FFT is applied to turn the data in Step 3 into a frequency plot, which is a static multidimensional representation (Step 5) of the shape in Step 1. Step 6: A scalar value can be created from Step 5 via an index. See the **Figure 4** for a discussion on types of indices to calculate. (**B**) A flowchart showing a simplified description of the steps of the LCPC transform for the parallel grid system. (**C**) Examples shape themes for which the parallel grid system is optimal.

While the radial grid system measures the distance of each intersection from the origin of the radial grid (**Figure 1A**), the parallel grid system always measures the distance of intersections with reference to an imaginary line at the left of the shape (**Figure 2A**). The open-source script determines the position of this imaginary line by finding the left-most pixel of the contour and then moving 10 pixels to the left of this position. Here, the x-coordinate of this position becomes the reference line from which all intersections are calculated. This 10-pixel rule is arbitrary, but it is why all contours analyzed by the open-source script for the parallel grid system must have at least 15 pixels of white space on all four sides of a contour.

For 2D shapes that have folds or multiple layers, gridlines may intersect with the contour more than once. In this case, the distances of all intersections on a gridline are summed into one value. Thus, each gridline has only one x-coordinate and one y-coordinate. This summation is represented by the term “compressed” in the name LCPC transform. The rendering of a non-linear 2D shape into a discrete sinusoid wave is represented by the term “linearized” in the algorithm’s name. Lastly, the term “polar coordinates” is in the algorithm’s name because the first grid system conceptualized was a 180-degree radial grid of polar coordinates^7^. Even after the realization that polar coordinates and Cartesian coordinates are interchangeable, the name of the algorithm was kept as is. The acknowledgements section describes the personal reasons that motivated the invention and augmentation of the LCPC transform.

Protocol 1 was the sequence for obtaining the data shown in **Figure 3B**, while Protocol 2 was the sequence for obtaining the data shown in **Figure 3C-3D**. The steps in these protocols have individual Python scripts that are available in the GitHub repository called “Pre-Processing-Tools-for-LCPC-Transform”^16^. Protocol 1 is an example of the pre-processing steps to derive “pure shape” in preparation for the LCPC transform. Protocol 2 is an example of pre-processing steps to measure shapes “at scale”, meaning at their original scales relative to each other. For **Figure 3**, tissue specimen were collected at Brigham and Women’s Hospital and the University of California, San Francisco under Institutional Review Board (IRB)-approved protocols from patients who provided informed consent for research use of their tissues, as previously described^20^.

**Figure 3:**
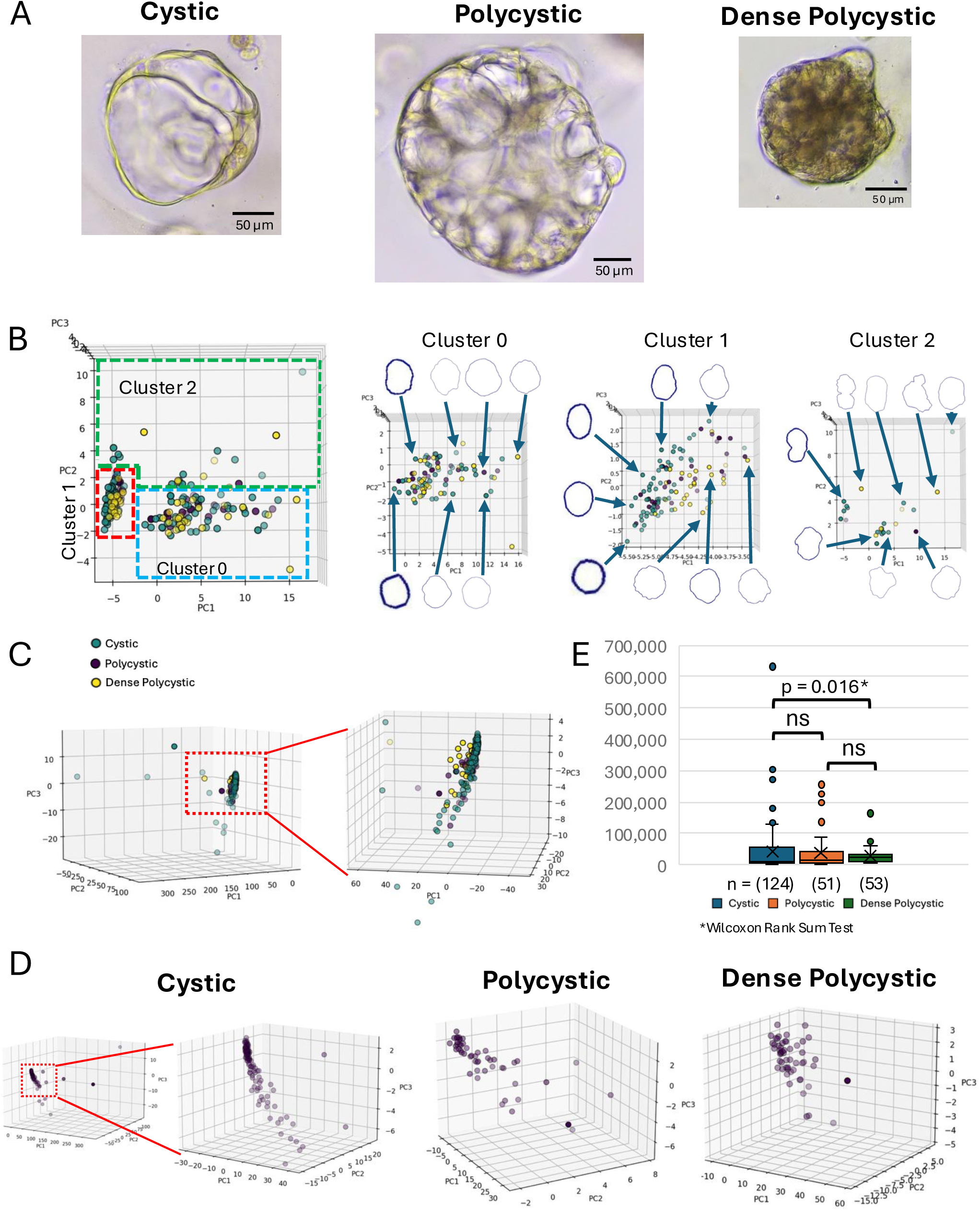
The LCPC transform objectively identifies morphological subtypes of breast cancer organoids. (**A**) Primary human breast organoids were derived as previously reported^19–21^ and as described in **Supplementary File 1**. Three phenotypes could be observed and classified based on visual inspection: cystic, polycystic, and dense polycystic. Scale bar = 50 µm. (**B**) The outer contours of the organoids were manually segmented (**Supplementary Figure 1**; criteria described in **Supplementary File 1**) and the pure shape of these contours was obtained. Note that the contours in the diagram are all rotated such that the longest intern line is vertical, thus do not match their original orientations. The radial grid LCPC transform was then applied, and the results were analyzed via PCA. K-means Clustering identified three clusters (0, 1, and 2), and Silhouette Scores (SS) were calculated to confirm the existence of these clusters. Cluster 0 vs 1, SS: 0.68; Cluster 0 vs 2, SS: 0.27; Cluster 1 vs 2, SS: 0.69. The contours on the dot plot were oriented such that the longest internal line is vertical; thus, not the original orientation of the organoids. (**C**) The outer contours of the organoids were also measured via the radial grid LCPC transform on their at-scale shapes, meaning the shapes as they are in the images relative to each other without any resizing. The inset is enlarged to show the looser packing of polycystic and dense polycystic organoids compared to the cystic organoids. For B and C: Green = cystic, Purple = polycystic, Yellow = dense polycystic. (**D**) The first three PC values are plotted for each phenotype separately and the average KNND is calculated to show that the cystic phenotype is most densely packed, while the polycystic and dense polycystic phenotypes exhibit less density (meaning more heterogeneity). Though plotted separately, all three phenotypes share the same PCA space. For the cystic phenotype, average KNND was calculated on the inset because the outliers that exist only in this phenotype skew the calculation. Average KNNDs: Cystic, 1.061; Polycystic, 1.575; Dense Polycystic, 2.247.

Segmentation can be done manually in image processing software, such as Mac’s Preview software or Microsoft’s Paint, or thresholding-based methods. If done manually, one of the following four colors should be chosen: blue, green, pink/magenta, red. The GitHub repository called “Pre-Processing-Tools-for-LCPC-Transform”^16^ contains a folder called “color extraction scripts”. This folder contains four Jupyter Notebook files that have Python code that extracts the four aforementioned colors and turns them into blue masks on a white background. This process can also be done via thresholding for the outline color in an image processing software such as Fiji/ImageJ. For images that contain contours that have touching borders or overlapping edges (like a Venn Diagram), they will need to be separated via an image processing software such that they become independent objects in the composite mask image that contains multiple organoid contours. This does not apply to images that only have one organoid or that have multiple organoids that don’t touch each other.

## PROTOCOL

### 1. Extracting pure shape of 2D organoid contours for the LCPC transform

1.1. Segment the outer edge of each organoid using any image processing software, such as Microsoft’s Paint, MacOS’ Preview, or Fiji/ImageJ.

1.2. Extract the contours of the organoids as open masks that are blue lines on a white background.

1.3. Isolate each blue contour onto its own white canvas.

NOTE: This step is only necessary if an image contains more than one contour.

1.4. Crop each contour to have a one-pixel margin on all four sides of the blue object.

1.5. Add 100 pixels of white space on all four sides to create new margins.

1.6. Rotate the objects such that the longest internal length is horizontal.

NOTE: This step turns the longest internal line with each object into the object’s width. There are two scripts available for this purpose. For closed shapes, use the Jupyter Notebook named “Rotate CLOSED Object Horizontal by Longest Internal Length_v2.ipynb”. For open shapes or shapes with multiple components, use the Jupyter Notebook named “Rotate Green Line_v3.ipynb”. The Rotate Green Line tool has a video tutorial that is linked in the ReadMe file contained within the same folder as the .ipynb file.

1.7. Crop each image to have a one-pixel margin on all four sides of the blue object.

NOTE: This is the same as Step 1.4. The purpose of trimming this time, however, is that the resizing step that comes next resizes the entire canvas, not just the blue object in it. Thus, by making the blue object nearly the same width and height as the canvas itself, resizing the canvas to 400 pixels wide also resizes the object to be nearly 400 pixels wide. There is a Jupyter Notebook named “Trim margin to 1-pixel border.ipynb”.

1.8. Resize the canvas width of each image to be 400 pixels wide while also constraining the aspect ratio.

NOTE: This step makes each object have the same width. Constraining the aspect ratio prevents skewing the object during the resizing. There is a Jupyter Notebook named “Resize width to 400 pixels but constrain aspect ratio.ipynb”.

1.9. Add 100 pixels of white space on all four sides to create new margins.

1.10. Rotate each image either 90 degrees clockwise or counterclockwise to make the longest internal length vertical, being consistent with the choice for all images.

NOTE: In the above sequence, pure shape is obtained by sequentially implementing Steps 1.6, 1.7, 1.8, and 1.9. Adding margin space in Step 1.9 is required to avoid errors in the LCPC transform scripts that will be done next, but is not involved in extracting pure shape. Users who desire to measure shapes “at-scale” need not perform Steps 1.6, 1.7, and 1.8 in the above sequence, though they should still pre-process their mask images to attain optimal orientation prior to applying the LCPC transform.

1.11. Perform the LCPC transform on each contour. Use Python scripts provided for the radial grid method^17^ or the parallel grid method^18^.

### 2. Extracting at-scale shape of 2D organoid contours for the LCPC transform

NOTE: The step-by-step method for extracting the at-scale shape of objects is like that for extracting pure shape in Protocol 1 above. The steps are named identically in both protocols for ease of matching. Please refer to the notes in Protocol 1 for each step. The main difference between pure shape and at-scale shape is that at-scale shape does not require Steps 1.6, 1.7, and 1.8, which is the sequence for resizing objects.

2.1. Segment the outer edge of each organoid. Use a computational approach or do it manually via a basic image editing software, such as Microsoft’s Paint, MacOS’ Preview, or Fiji/ImageJ.

2.2. Extract the contours of the organoids as open masks that are blue lines on a white canvas. Use Fiji/ImageJ to extract masks or the Python script provided^16^.

2.3. Isolate each blue contour onto its own white canvas. Use Fiji/ImageJ to extract masks or the Python script provided^16^.

2.4. Crop each contour to have a one-pixel margin on all four sides of the blue object. Use Fiji/ImageJ to extract masks or the Python script provided^16^.

2.5. Add 100 pixels of white space on all four sides to create new margins. Use Fiji/ImageJ to extract masks or the Python script provided^16^.

2.6 Rotate each contour such that its longest internal line is vertical. Use Fiji/ImageJ to extract masks or the Python script provided^16^.

2.7. Perform the LCPC transform on each contour. Use Python scripts provided for the radial grid method^17^ or the parallel grid method^18^.

## REPRESENTATIVE RESULTS

### The radial grid LCPC transform applied to breast cancer organoids

3D organoids and tumors can adopt a variety of shapes, some of which are obviously different to the human eye, even though traditional metrics yield statistically insignificant differences. On the other hand, organoids can also exhibit shapes that seem heterogeneous, and thus insignificant, to the human eye, masking recurring subtle morphologies that represent distinct subtypes. Human primary breast organoids were extracted from patients and cultured in 3D following a previously published protocol^19,20,21^. **Figure 3A** shows three morphological categories accessible based on visual inspection: a cystic phenotype, a polycystic phenotype, and a dense polycystic phenotype. **Figure 3B** shows that the outer contours of these organoids were manually segmented and their pure shape was extracted, which removes the effect of size on their shapes. Because the contours were circular in nature, the radial grid LCPC transform was applied to the pure shapes followed by PCA. Note that the contours overlayed on **Figure 3B** were rotated such that the longest internal line is vertical, meaning their orientations are not the same as in their source images. **Figure 3B** shows a dot plot of the first two principle components (PCs) and that K-means clustering identified three clusters: 0, 1, and 2. Silhouette Scores (SS) confirmed the likely existence of two separate clusters between Clusters 0 vs. 1 (SS = 0.68) and Clusters 1 vs. 2 (SS = 0.69), while indicating a weak separation between Clusters 0 vs 2 (SS = 0.27). Despite the weak Silhouette Score between Clusters 0 and 2, labeling the data points with the pure shape of each data point justifies that Cluster 2 represents the most distinct phenotype: highly asymmetrical, non-circular contours (the thickness of the contours does not matter in this case).

While pure shape removes the effect of size on shape, which is impossible when measuring area and volume, measuring the organoid contours at-scale also reveals interesting insights. **Figure 3C** shows a 3D dot plot of the first three PCs resulting from the radial grid LCPC transform performed on contours as they were segmented from the images without any resizing. This approach means that the LCPC transform is measuring both the shape and size of the organoids simultaneously. **Figure 3C** shows that the polycystic and dense polycystic organoids are less densely packed compared to the cystic organoids. To quantify this expansion, **Figure 3D** shows the scatter of each phenotype in their shared PCA space. The average K-nearest Neighbor Distance (KNND) is calculated for each phenotype, which supports the visual assessment of decreasing density of the dots going from cystic organoids (KNND = 1.061) to polycystic organoids (KNND = 1.575) to dense polycystic organoids (KNND = 2.247); this decreased density of dots can also be interpreted as increased morphological heterogeneity of outer contours. Lastly, for a comparison of the LCPC transform to the traditional metric of area, **Figure 3E** shows that while there is a statistical significance between the area of cystic organoids vs. dense polycystic organoids (p = 0.016, Wilcoxon Rank-Sum Test), there is no indication based on area that there are three distinct morphological groups across the phenotypes as identified by K-means Clustering after pure shape LCPC analysis (**Figure 3B**) or that there is increasing morphological heterogeneity as show by at-scale LCPC analysis (**Figure 3C–D**). **Figure 3E** shows a decreasing scatter of the data points going from cystic to polycystic to dense polycystic organoids, which is the opposite of what is observed by at-scale LCPC analysis (**Figure 3C**), highlighting that the spatial information captured by the LCPC transform is different from what area captures.

The same approach applied to individual 2D shapes in **Figure 3** and **Supplementary Figure 2** also apply to the contour of cells in 2D cell culture: single-cells (such as red blood cells) or clusters of cells (such as in cervical Pap smears). Pilot studies for each of these cases are described on YouTube: shape analysis of brain organoids that exhibit varying degrees of folding in their outer contour^22^; shape analysis of lung alveoli *in situ*, the same principles can be applied to analyzing 2D cell culture^23^; shape analysis of cervical tissue from Pap smears in which both the contour of the nucleus and of the cell membrane are simultaneously quantified via the radial grid LCPC transform^24^; shape analysis of red blood cells infected by malaria and cells adjacent to infected cells but are not themselves infected^25^; shape analysis of red blood cells affected by Sickle Cell Disease^26^.

### The parallel grid system for measuring density or texture as an index of spatial information

Spatial information involves more than the usual notions of shape, such as squareness, roundness, and curvature, because it encompasses concepts such as texture and density. Complex tubular networks, such as blood vessels, or branching networks, such as cytoskeletal filaments, do not fit the notion of shape in the same sense of our geometry-based intuition of shapes, such as circles and polygons. However, density and texture are very useful notions of spatial information for understanding the structure and function of cells, organoids, and tumors.

In dealing with how to quantify complex networks in fluorescence imaging, cell biologists measure features such as the average signal intensity within an image or the percentage of the total area of an image that contains a signal. These approaches recognize that there is more to spatial information than what is described in terms of geometry motifs.

The pattern of vascular networks within tissues, tumors, and organs on a chip is not easily defined by traditional geometry motifs, but benefits from the interpretation as texture and density. Vascular remodeling in tumor development and treatment resistance is a well-established feature of cancer (reviewed in^27^). Histopathological sections of 458 primary neuroblastic tumors were analyzed^28^, characterizing the density, size, and shape of total blood vessels and vascular segments. They found that blood vessels were larger, more abundant, and more irregularly-shaped in tumors of patients with poor prognostic factors compared to the favorable cohort. Interestingly, a term called “vessels that encapsulated tumor clusters” (VETC)^29^, was coined to describe a pattern of blood vessels that surround clusters of small tumors within a larger hepatocellular carcinoma and are associated with higher rates of metastasis and recurrence. The same group that coined the term later founded that VETC-positive hepatocellular carcinoma correlated with a positive response to treatment with the kinase inhibitor Sorafenib^30^. Thus, the ability to quantify vascular patterns is crucial for prognosis and understanding mechanisms of resistance.

For quantifying dense networks of tubes, islands, and branches, the parallel grid system of the LCPC transform is useful for measuring what can be called texture-density, instead of shape. As an example of how the parallel grid LCPC transform can do this, **Figure 4A** shows cardiac microvasculature patterns from a study of cardiac vasculature^31^: to the naked eye, objectively and quantitatively distinguishing the patterns between the infarcted areas and the areas at a region remote from the site in injury is very difficult. By extracting the mask of the blood vessels (**Figure 4A** Step 1) and applying the parallel grid LCPC transform, quantitative differences can be observed. First, to enhance the cleanliness of the data, each image was cropped with a circle of uniform diameter, and that is centered at the middle of the square image (**Figure 4A** Step 2), which ensures that each image has the same height and width regardless of rotations. Next, the cropped images are rotated such that the main axis of vessels (red double arrows in **Figure 4A** Step 1) is vertical (**Figure 4A** Step 3). Then, a blue line is applied to the circumference of the image to ensure that even images with sparse vascular patterns have the same diameter in terms of blue pixels (**Figure 4A** Step 4). Steps 3 and 4 were performed using the Fiji/ImageJ software (v2.14.0). Lastly, the parallel grid LCPC transform is applied (**Figure 4A** Step 5). Analyzing the results via PCA and plotting the first three PCs reveals that the infarcted regions cluster separately from the remote regions (**Figure 4B**). Calculations of Chamfer Distance (CD) quantify the visual patterns exhibited by the graphs. At 1-day after injury, the infarcted site exhibits a vascular pattern that is more distant from the basal condition (Basal vs. 1Day_infarct: CD = 83.89) than is the non-injured site (Basal vs 1Day_remote: CD = 53.95). At 1-day after injury, the non-injured remote site is closer to the injured site (1Day_remote vs 1Day_infarcxt: CD = 34.48) than it is to the basal condition. At 3-days after injury, the vascular patterns of the injured site and associated non-injured remote site exhibit drastic deviation from the basal condition and from each other (Basal vs. 3Days_Infarct: CD = 240.63; Basal vs. 3Days_Remote: CD = 67.28; 3Days_Remote vs. 3Days_Infarct: CD = 168.35). Very interestingly, at 7-days after injury, the vascular patterns of the injured site and the non-injured remote site occupy a 3D space that is approximately orthogonal to the basal condition, while aligned with each other at a distance that is closer than they were at 1-day post injury (CD = 24.76 for day 7 compared to 34.48 for day 1). The data suggest two things: (1) as healing progresses, the vascular pattern of the injured site and non-injured remote site adopt patterns that are distinct from the basal condition; (2) the reconstruction program activated at the injured site also triggers similar morphological changes at their associated non-injured remote sites.

**Figure 4:**
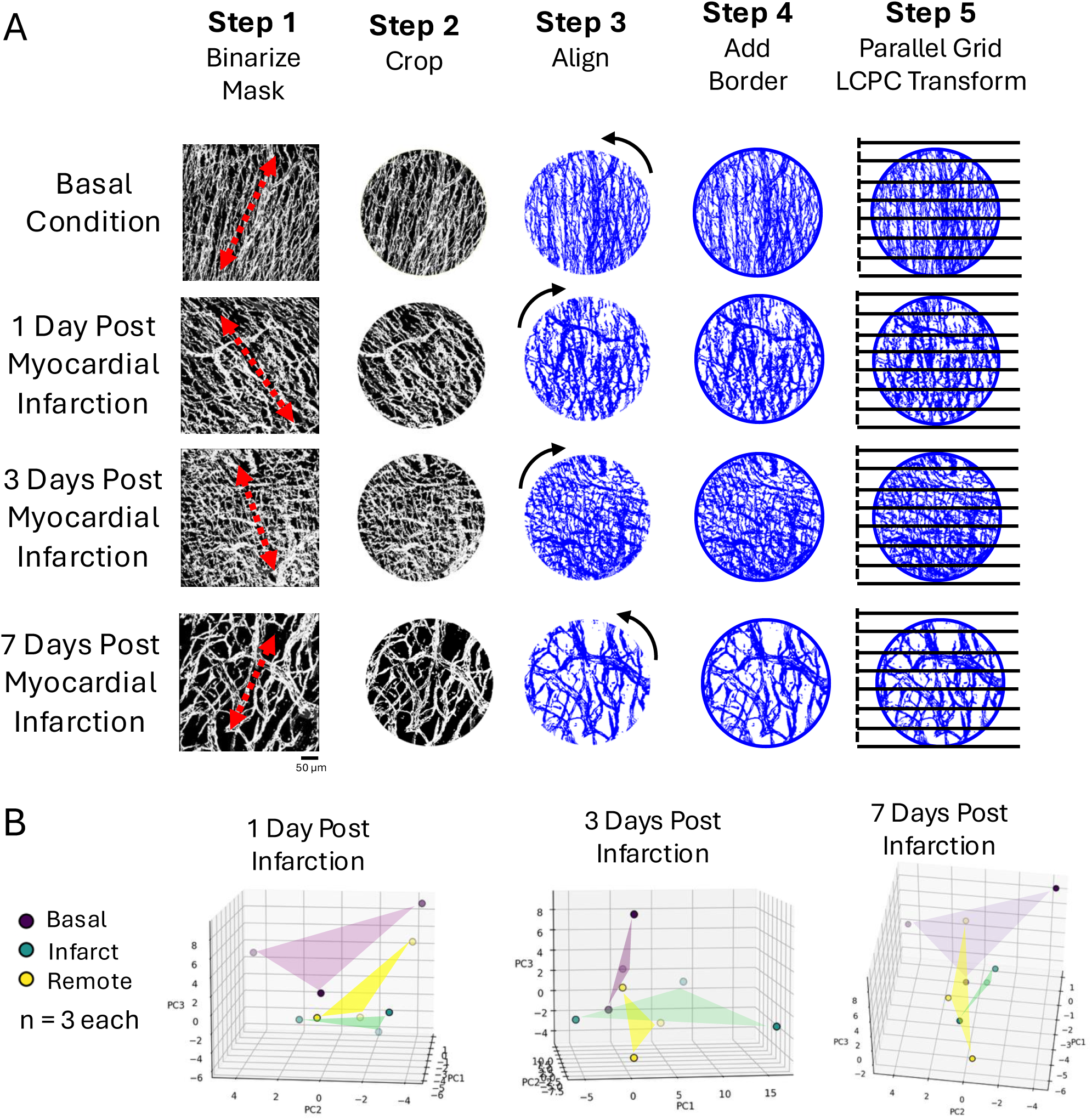
Measuring texture-density with the parallel grid LCPC transform. (**A**) Photomicrographs of microvasculature in cardiac tissue previously obtained^31,32^. Images from the remote sites that were non-injured in each time point are not shown. Step 1: A 2D mask is generated and the general axis of alignment of vessels is determined (red double-arrow). Step 2: A circular region centered on the midpoint of the image is obtained in order to force each image to have the same height after the images are rotated to varying degrees in the next step (rotating square images to varying degrees will alter the height of the image depending on the degree of rotation). Step 3: Each image is rotated such that the general axis of alignment determined in Step 1 is vertical. Step 4: A uniform blue border is applied around each image to ensure that each image has the same diameter, because images with sparser densities can be shorter (example: see “7 Days Post Myocardial Infarction” in Step 3). Step 5: The parallel grid LCPC transform is applied. Scale bar = 50 µm. (**B**) Dot plots of the first three PCs of the output from the parallel grid LCPC transform (Purple: Basal condition, Green: infarcted site, Yellow: non-injured site remote to the infarcted site). See the results section for Chamfer Distance calculations at each time point between the three conditions.

As a reference for the versatility of the LCPC transform, the traditional metrics of percent area (**Figure 5A**) and total perimeter (**Figure 5B**) of blood vessels was calculated. While area and perimeter reveal changes between points in the time course, they do not provide coherent quantitation of distinct spatial arrangements of vessels. Area and perimeter are abstractions that can represent many different vascular arrangements and thus are not informative about how the spatial arrangement of vessels are changing across the treatment conditions and time course. With an n = 3 for each group, the t-test and Wilcoxon test are unreliable. Cliff’s Delta is nonparametric and works well to show a shift in the effect size. The major difference revealed by measuring percent area of blood vessels is between the basal condition and the 7-day time points; for total perimeter of blood vessels, the differences are between the basal condition vs. 1-day and the basal condition vs. 7-days. However, area and perimeter provide no indication that there are differences between the infarcted sites and their non-injured remote sites, which is very surprising given that the infarcted site was purposefully injured. Here is where the versatility of the LCPC transform emerges compared to traditional metrics. For each sample, the results of the FFT as part of the LCPC transform produces a set of magnitudes associated with a set of frequencies. The magnitudes of each frequency bin can be averaged and then compared across the treatment groups (**Supplementary Figure 2**). This allows for a simple assessment of which frequency bins exhibit the biggest differences between treatment groups. **Figure 5C** shows six bins that exhibit clear effect size differences between various treatment groups, especially between the infarcted sites and their associated non-injured remote sites, which area and perimeter indicated had no difference. The independent values in these six bins provide multiple indices that show quantitative differences between treatment conditions. Interestingly, the pattern between the groups for Bin 1 of the LCPC transform results is similar to the result obtained from measuring total perimeter. The other indices, however, reveal quantitative differences that are hidden in both area and perimeter. Indices unique to the basal condition: Bin 49, Bin 56, Bin 21. Indices unique between the Day 1 groups: Bin 1, Bin 6, Bin 49. Indices unique to the Day 3 groups: Bin 6, Bin 112, Bin 49. Indices unique to the Day 7 groups: Bin 6, Bin 49, Bin 112. These data highlight the versatility and precision of the LCPC transform over the traditional approaches of area and perimeter.

**Figure 5:**
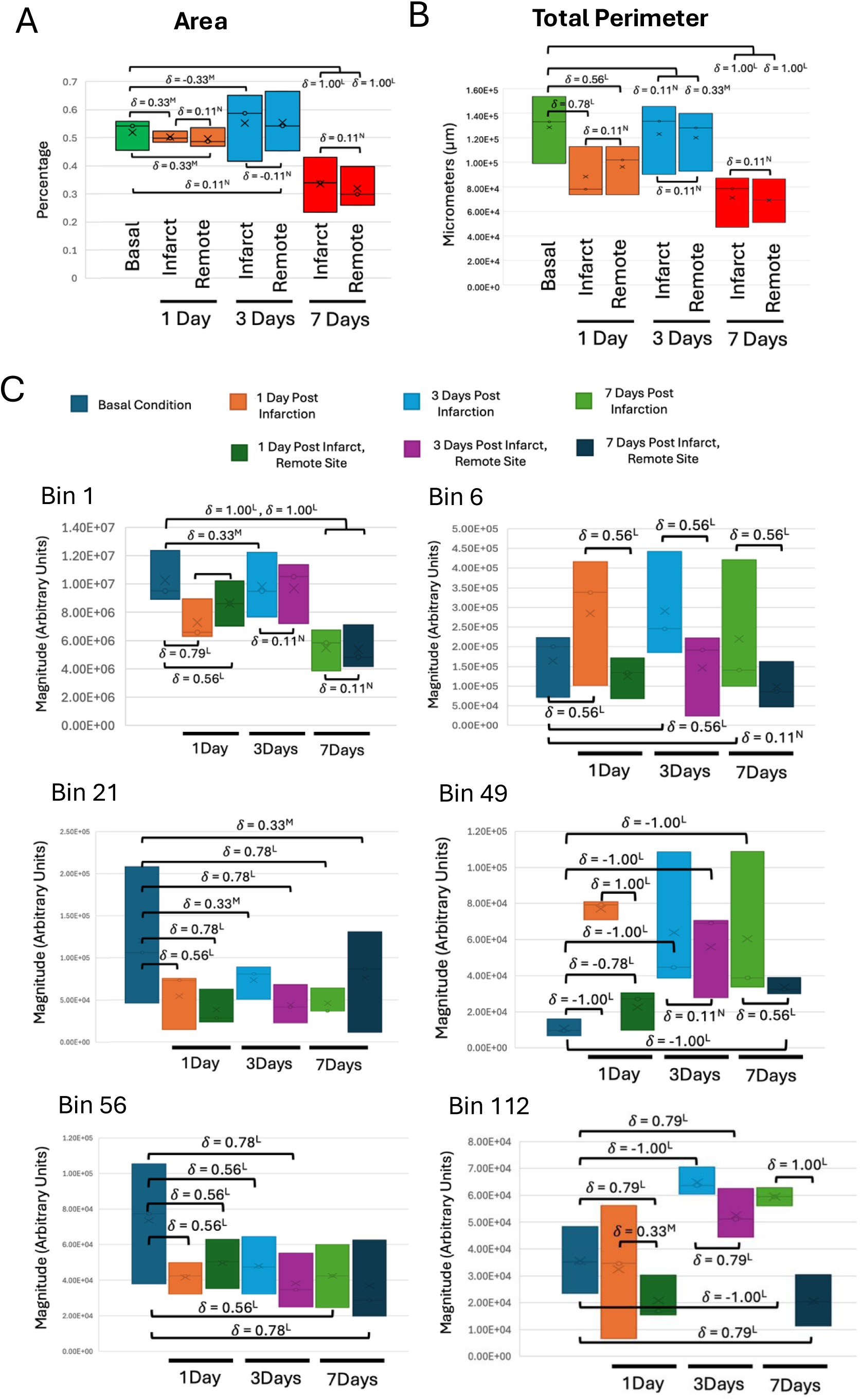
The LCPC transform measures features hidden to area and perimeter. For a comparison to traditional shape metrics, the percent area (**A**) and the total perimeter (**B**) of the vascular networks were measured. (**C**) To show the precision that can be obtained from the result of the LCPC transform, the value of individual frequency bins resulting from the FFT were compared (**Supplementary Figure 1**). Six frequency bins were chosen as indices that represent the objective quantitation of differences between treatment groups. For all panels, due to the limited sample size of each group (n = 3), the nonparametric Cliff’s Delta for effect size was calculated instead of a p-value from the Wilcoxon Rank-Sum Test.

## DISCUSSION

As with any computational tool, the quality of the output depends on whether the inputs adhere to the expected rules around which the tool was designed. The following rules should be followed and used as quality control checks when utilizing the provided Python scripts. First, make sure that that open masks are blue lines on a white background. Blue was an arbitrary decision and has no significance, but the scripts that perform the LCPC transform were written to search for blue pixels on a white background. Second, do not run the LCPC transform scripts on images that have less than 10-pixels of white space on all four sides. See this video for an explanation of why: “The Margin Thickness Parameter - How to Prepare Shape Outlines”^33^. Third, this script assumes that the outlines of your object are blue pixels on a white background. The thickness of the line should be between 5-10 pixels thick. See this video for an explanation of why: “The Line Thickness Parameter - How to Prepare Shape Outlines”^34^. Fourth, do not run the LCPC transform scripts on blank images. This will cause an error in the script. Fifth, do not run the LCPC transform scripts on images that consist of only a speck of debris. This will cause an error because the grid system may not detect the speck and thus attempt to analyze a blank image. Sixth, do not run the LCPC transform scripts on images that contain specks of unwanted debris. The debris will alter the height and width of the open mask object and thus cause inaccurate results. Seventh, see this video to understand why it is important to understand the scale of your objects before analyzing them with the LCPC transform: “The Scale Parameter - How to Prepare Shape Outlines”^35^.

One of most amazing aspects of the LCPC transform is its ability to capture spatial context. Biology is full of spatial context that is crucial for understanding nature. For example, epithelial cells have apical-basal polarity^36,37,38,39^. Furthermore, the direction of gravity, which is altered in microgravity conditions, has significant effects on both astronaut physiology^40,41^ and cancer cell morphology^42,43,44^. Lastly, in situ pre-neoplasia and neoplasia can grow into lumens via a stalk that serves as the central support structure that houses access to the blood supply. So, when these lesions regress, they regress both away from the luminal cavity and towards the stalk^45,46^. Because it was designed to do so, the LCPC transform is highly sensitive to spatial context and thus can introduce extra heterogeneity if the user is not careful. Figure 6 describes how to systematically orient shapes before applying the LCPC transform. Figure 7 describes how traditional geometry cannot measure shape concepts that are obvious to humans. Figure 8 provides a diagram visualizing the difference between measuring pure shape vs. at-scale shape. Figure 9 describes spatial context is often overlooked when quantifying shapes in 2D culture, histology, and 2D slices from 3D imaging. Note that due to space limitations, **Supplementary File 1** contains extensive discussion of Figure 6 and Figure 7 in Section 5, and Figure 8 and Figure 9 in Section 6.

**Figure 6:**
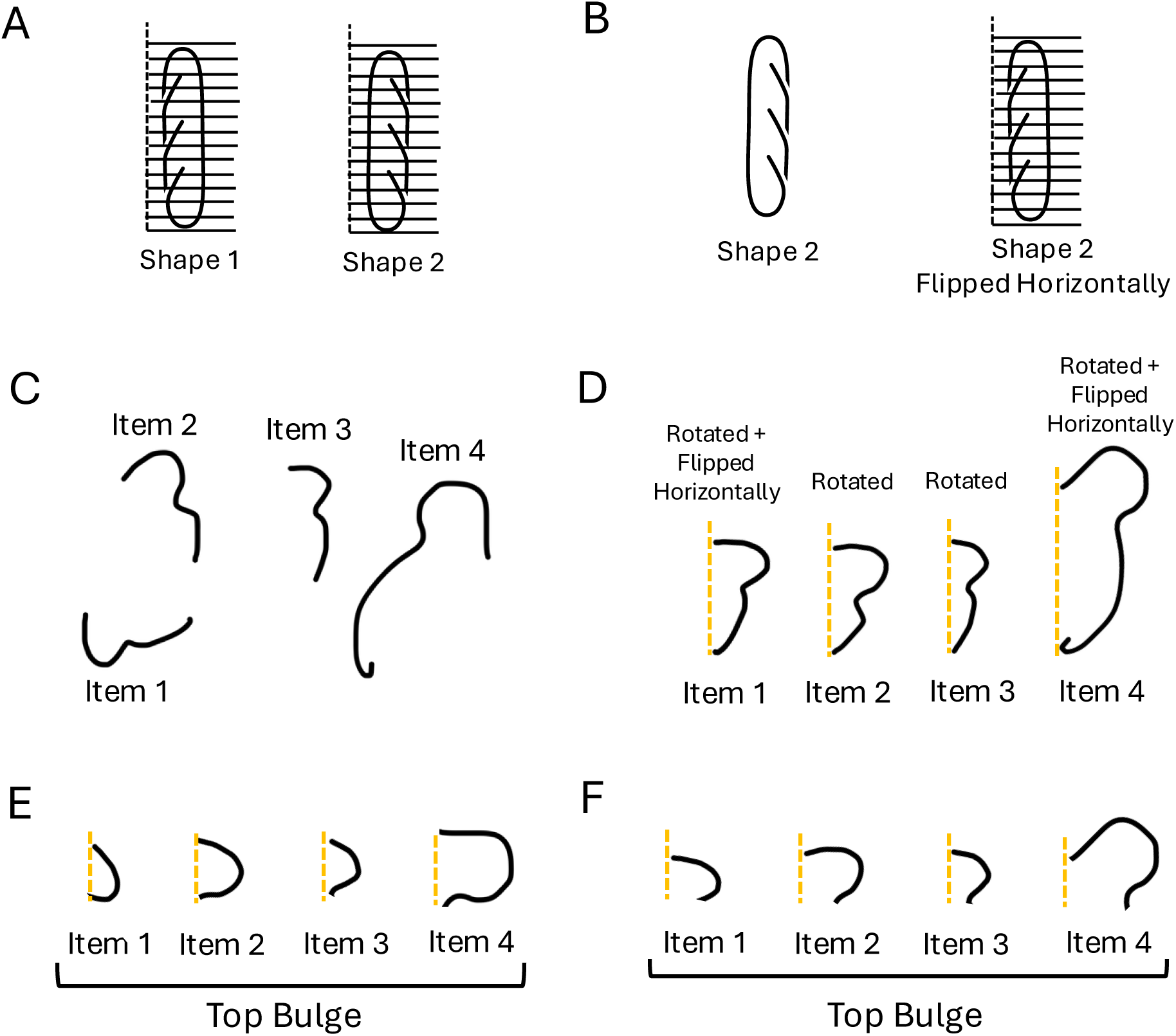
The importance of orienting shapes systematically for extracting clean data. (**A**) An example of identical shapes that are mirror images of each other and how the parallel grid system determines their distinct orientation and thus will produce different results. (**B**) An example of how to flip Shape 2 horizontally, such that it’s orientation now matches Shape 1 and becomes in a fairer comparison to Shape 1. In this configuration, the parallel grid LCPC transform will yield the same results for both Shape 1 and Shape 2. This alteration of orientation should only be done if there is no spatial context that suggests that Shape 2 should not be modified. (**C**) An example of four variations of objects that look like the number “3”, with some that face the wrong direction. (**D**) An example of how to orient each object to maximize the cleanliness of the data that will result from the parallel grid LCPC transform. (**E**) An example of measuring the wider bulge of the items in Panel C based on the same orientation rules as for Panel D. (**F**) An example of measuring the large bulbs of the items in Panel C but keeping their alignment context as in Panel D. The motivation of keeping the alignment rules in Panel D is to dissect the contribution of the spatial information in the wider bulge relative to the contribution of the narrower bulge within these rules. See **Supplementary Figure 3** for the FFT spectra associated with C-F. The images in A-B is an icon from Microsoft Powerpoint. The shapes in C-F were drawn in Microsoft Powerpoint.

**Figure 7:**
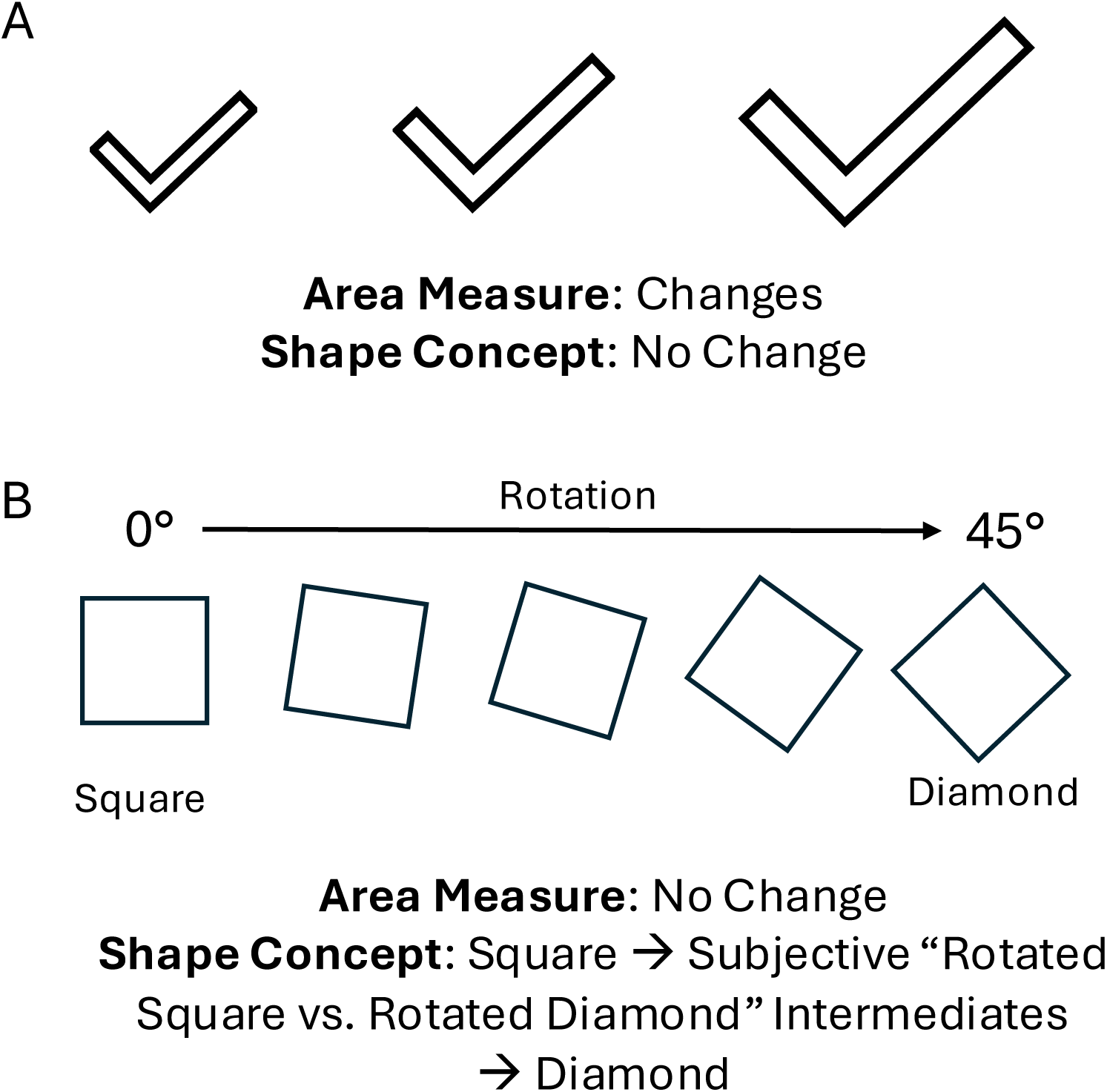
The inability of traditional shape metrics to register obvious context. (**A**) The series of checkmarks are the same shape but increase in size from left to right. In this series, the area of each checkmark is different while the shape concept remains the same: that is, they remain the same style of checkmarks. (**B**) A square is rotated about its center for 45-degrees until its shape concept becomes what humans call a diamond. While the shape concept of square is very distinct from that of a diamond, the in-between rotations are subjectively interpreted as either “tilted squares” or “tilted diamonds”. The LCPC transform can objectively and precisely measure the difference in each step in this series.

**Figure 8:**
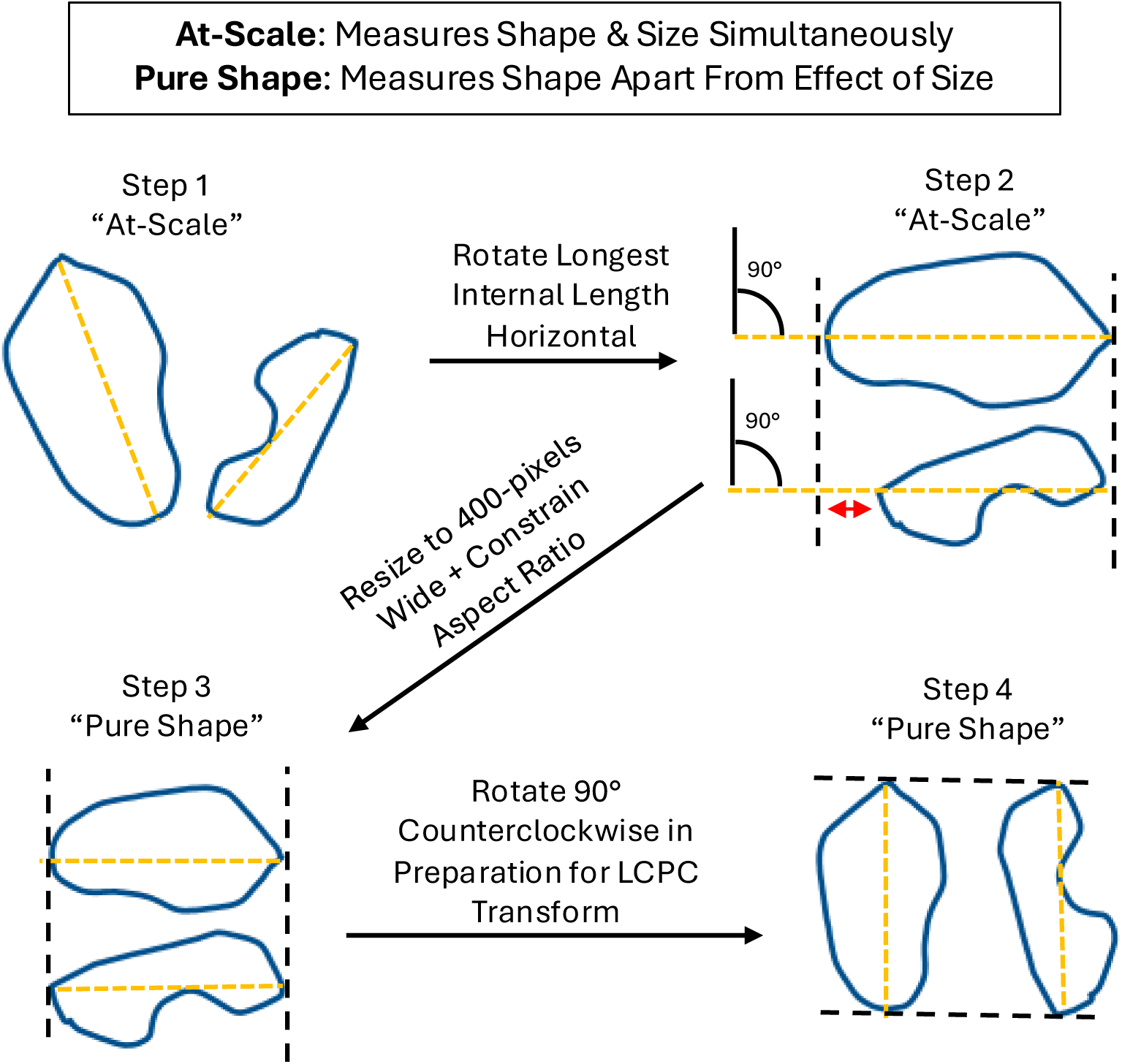
Explanation of pure shape vs. at-scale shape. Measuring objects “At-Scale” means measuring the shape of objects according to their original sizes relatively to each other. “Pure Shape”, on the other hand, means measuring the shape of different objects at a similar scale, which removes the effect of size on the shape. It is important to find an objective manner in which to resize the objects being compared. The longest internal line is often an optimal choice (Step 1). Both objects are rotated such that the longest internal line is horizontal (Step 2). In this case, the two objects are also rotated such that the indented side faces down. Next, the width of each object, which is now the same magnitude as the longest internal line of each object, is resized to be 400-pixels wide while also constraining the aspect ratio to prevent skewing the objects (Step 3). Lastly, it is optional to rotate the resized objects 90-degrees (Step 4) (in this case, counterclockwise) before applying the LCPC transform.

**Figure 9:**
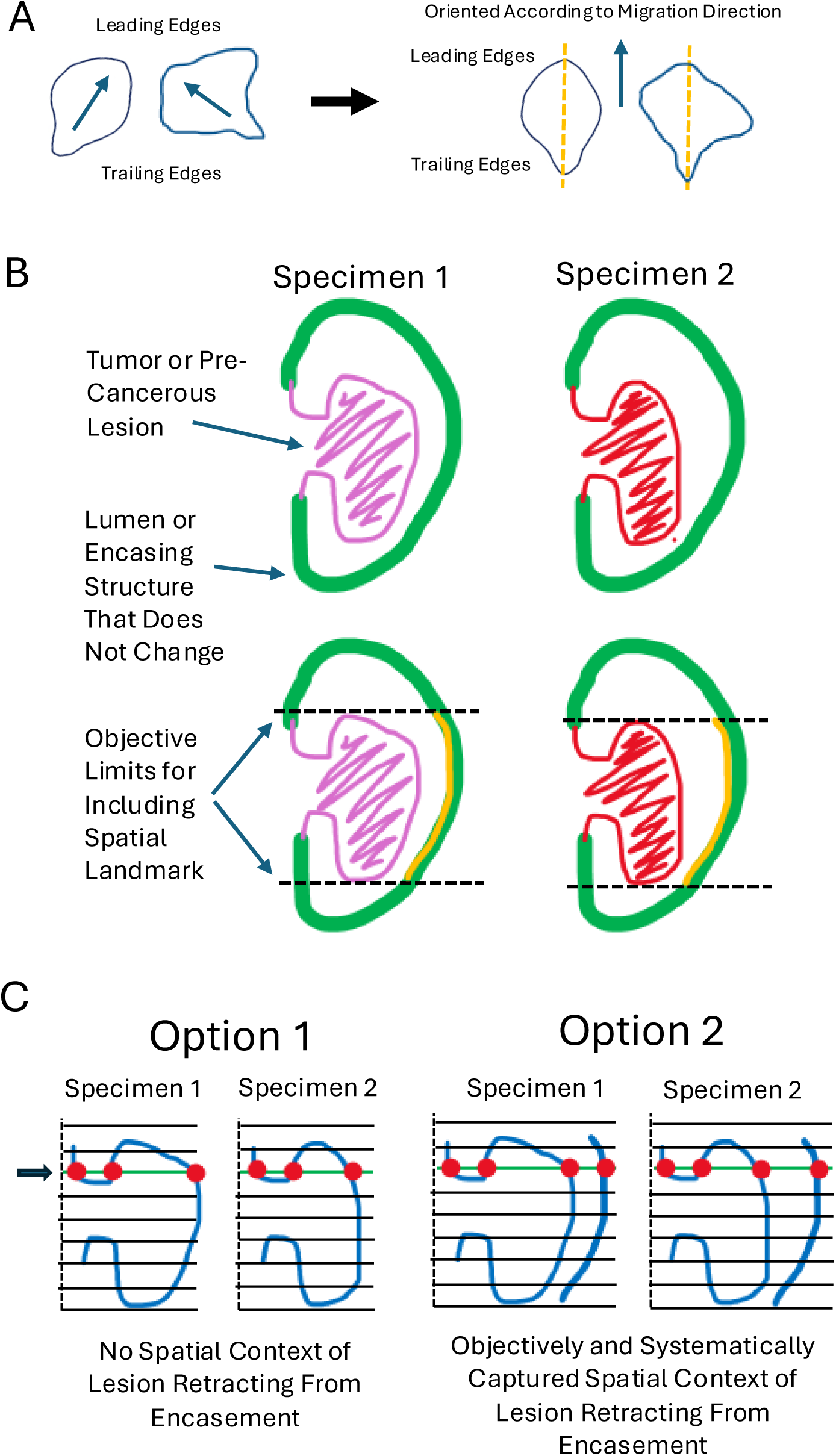
Example of spatial markers for cancer research. (**A**) A hypothetical example of cells migrating in 2D culture by expanding their lamellipodia in the direction of migration. In the context of migratory morphology of epithelial cells, orienting the contour of each cell such that the longest internal line that aligns with the axis of migration is vertical makes the most sense for capturing the spatial context. (**B**) Hypothetical examples of tumors or pre-cancerous lesions that grow within an encasement that does not change shape. The lesion grows from a stalk, which is the central support structure. As the lesion grows or shrivels, its contour moves towards or away from the internal wall of the encasement. The dotted lines represent a rational and objective way to define how much of the encasement should be traced (orange line) such that not too little or too much extra information is added. (**C**) Option 1 shows the parallel grid LCPC transform applied to the contour of the lesions without context of where the encasement is located. Option 2 shows the same but including spatial context of where the encasement is located. As an example, the red dots represent intersections between the grid line (black arrow, green line) and the contour. Option 2 results in each gridlines making a fourth intersection that does not exist in Option 1, which quantitatively adds spatial context to the result in Option 2 that is completely absent in Option 1.

Because the FFT results in multidimensional spectral data, it is important that the user become comfortable deriving indices from recurring patterns within the spectra. Though this manuscript refers to the LCPC transform as including the step of applying the FFT, the actual LCPC transform happens before FFT step (Figure 1A-B Step 4, Figure 2A-B Step 3). However, the effect of applying the FFT is to reveal objective, systematic insights from the frequency domain of the output of the LCPC transform. Thus, the LCPC transform would not be as effective without including the FFT as the final step. In the LCPC transform, the FFT results in a series of bins and their magnitudes, all of which are a static multidimensional representation of the contour inputted in Step 1. An analogy of a static multidimensional representation of information is the nine-digit phone number representing a person’s telephone communication channel in the United States. While cumbersome in comparison to scalar value representations of shapes (such as 85 cm in length, 56 degrees wide, etc.), multidimensional data harbors much more information for extracting insights. Figure 10 shows several spectral patterns that have been observed when applying the LCPC transform and how to derive indices from them. Section 4 of **Supplementay File 1** discusses Figure 10 more extensively.

**Figure 10:**
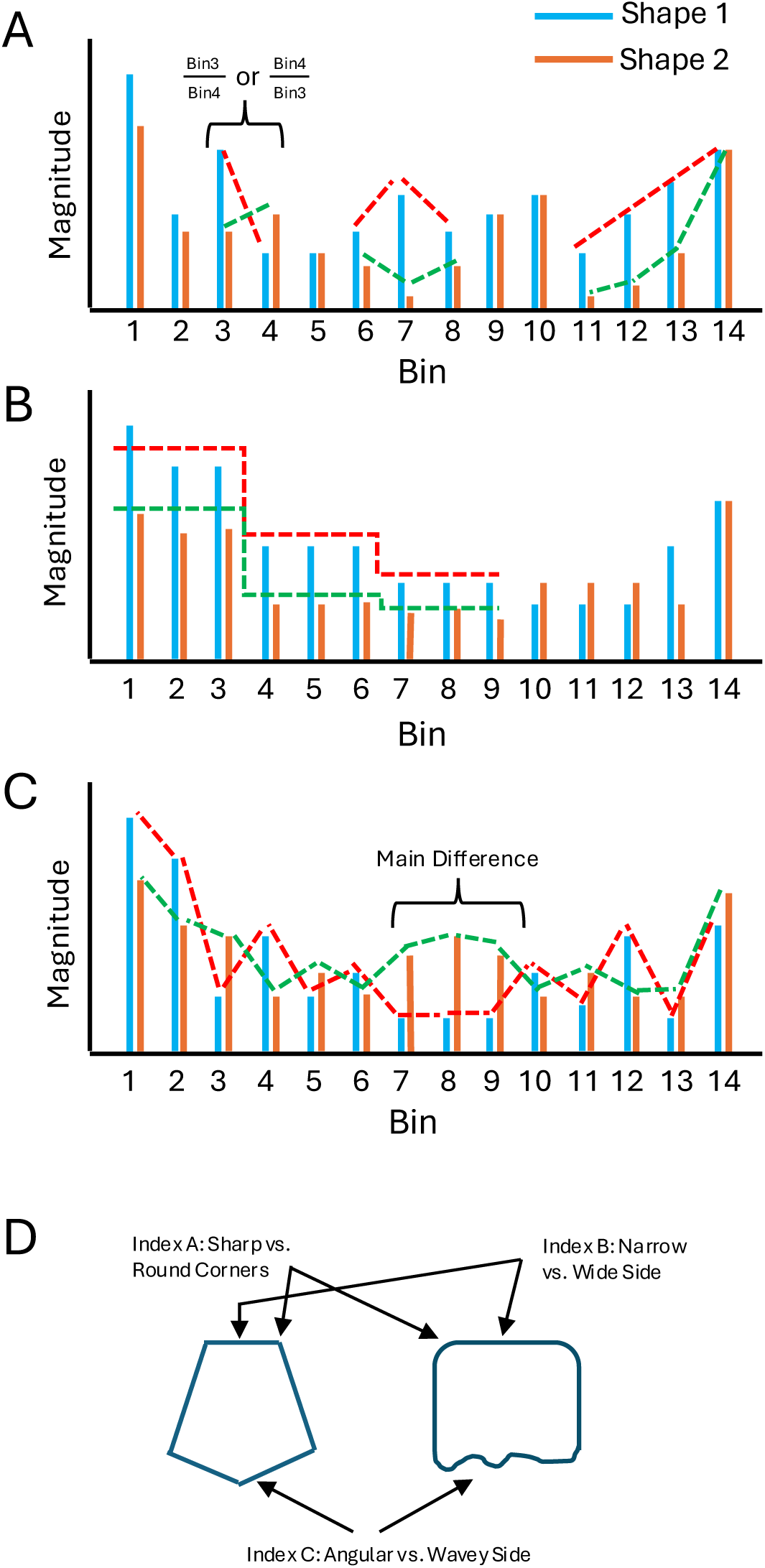
Hypothetical FFT results and common indices that can be derived from them. (**A**) The frequency plot resulting from the FFT of two hypothetical shapes is displayed. Since the magnitudes in Bin 3 and Bin 4 exhibit the opposite trend, a ratio between these two bins will be a good index representing a difference between these two shapes. The red and green dotted lines represent other frequency bins that could be used to derive good indices. (**B**) A histogram of two hypothetical shapes whose FFT profiles exhibit a stepwise decreasing trend as a representation of their morphological differences. (**C**) A histogram of two hypothetical shapes whose FFT profiles differ most in the central region along the span of frequency bins. (**D**) Examples of hypothetical morphological features that may yield indices that correlate with their presence.

The frequency domain of the LCPC transform reveals objective and quantitative features of a 2D shape that the naked eye cannot readily perceive. Very subtle differences in curvature, jaggedness, and tortuosity can be robustly quantified. However, once morphological subtypes are identified via this method, visually distinct themes can be observed that correlate with the quantitative differences. This feature makes the LCPC transform interpretable as a method of extracting spatial information for machine learning (ML) models and makes the resulting ML models themselves more interpretable. While neural networks are a powerful ML method, their results are often not interpretable to humans, which is a significant problem for the field of medical artificial intelligence (AI). Physicians are reluctant to employ an ML model for clinical decisions if they don’t understand why the model makes bad decisions. Furthermore, if no human understands why or how a model made a biased decision, then fixing this becomes intractable.

It is worth adding to the discussion of measuring texture-density (Figures 4 **and 5**) that the human eye is easily confused by optical illusions or disorderly patterns. Haphazard arrangement of complex patterns is often interpreted as being random, not measurable, or not worth measuring. The LCPC transform can measure reproducible, objective, quantitative differences in seemingly random patterns. However, unless the user is trained to not be misled by chaotic, disorderly patterns in nature that masquerade as randomness, no attempt will be made to measure the disorder to uncover predictable patterns. Predictable patterns are a clear indication that the system is not completely random. The LCPC transform was invented for the purpose of measuring shapes that are subtly different or inconceivably different to the naked eye. Because many morphologies in nature exhibit minor differences that seem more like meaningless chaotic heterogeneity than predictable subtypes of disorder – that each correlate with distinct biological functions – the following “Four Step” conceptual framework was developed to be used in conjunction with the LCPC transform. First, resist the notion that a complicated-looking system is random. Second, call the chaotic heterogeneity “disorder” and then find a type of order that can serve as a rational reference condition. Hypothesize that the disorder is deviating from the reference. Third, find a way of objectively measuring the pattern that is the reference, such as the LCPC transform. Fourth, apply this same objective method to measuring what is considered the disorder in system.

## ACKNOWLEDGMENTS

We would like to acknowledge Duane Nichols, a high school science teacher who passed away from colon cancer. This inspired the invention of the LCPC transform to characterize the morphology of colon polyps. Second, we would also like to thank Thuan Trinh, who suffered from Bipolar Disorder 2 and inspired the enhancement of the LCPC transform, such as adding markers to capture spatial context. The LCPC transform is informally called the Nguyen-Nichols-Trinh (NNT) transform. Third, we would like to thank Paul Leal, the lead angel investor who supported the startup that attempted to commercialize the LCPC transform. Fourth, we would like to thank Polyxeni Gkontra, Ph.D., for sharing photomicrographs of cardiac vasculature for this study. Fifth, we would like to thank the DF/HCC Breast SPORE: Specialized Program of Research Excellence (SPORE) (NCI 1P50CA168504), the UCSF Breast Care Center surgical and pathology teams and the Breast Care Center interns for assistance with tissue specimen, and funding support from the National Institutes of Health (R01CA281361).

---

**Supplementary Figure 1:**
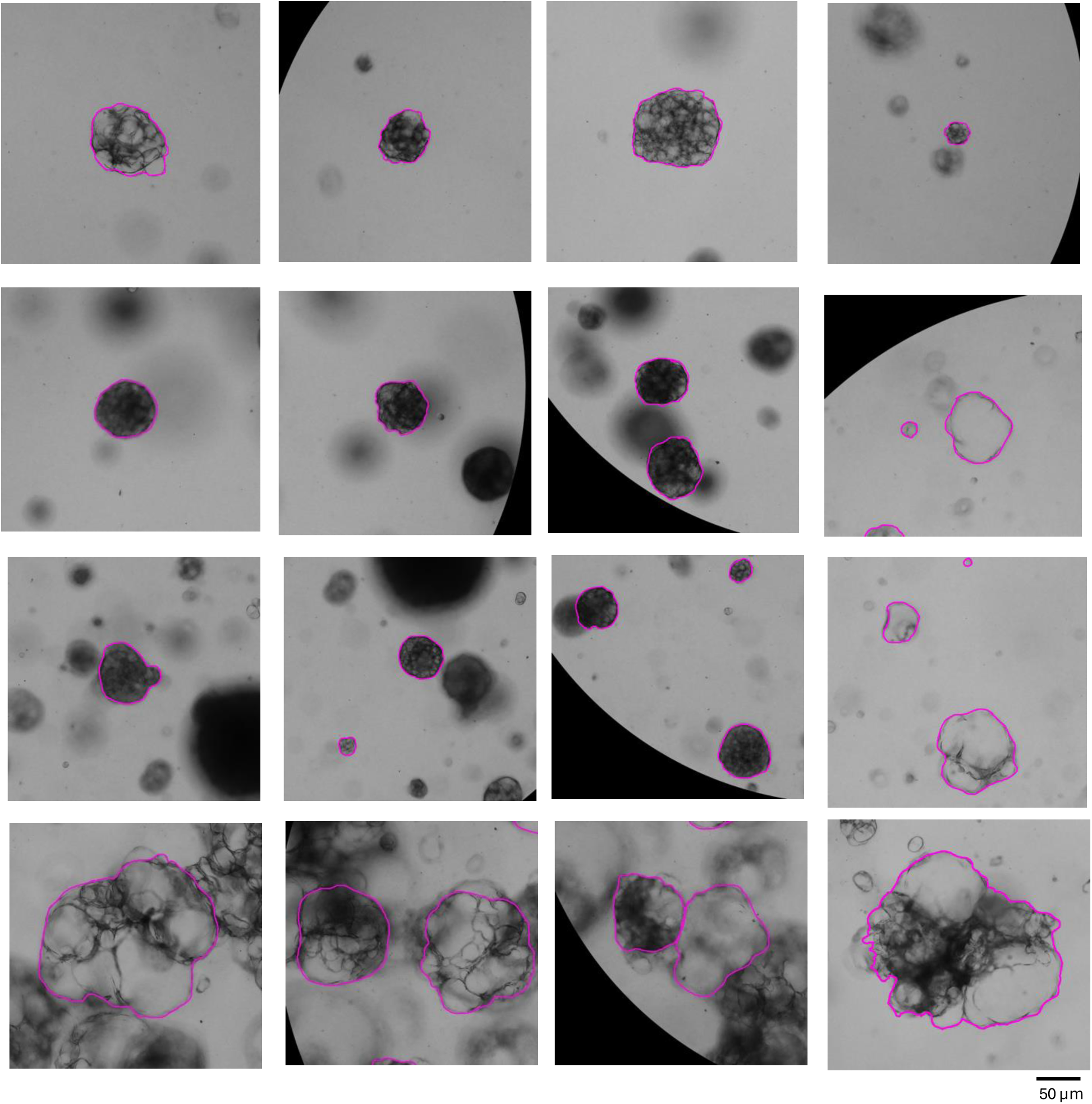
Example of segmented breast tumor organoids. Organoids were manually segmented by an expert who routinely cultures them and views them under microscopy. This figure displays examples of the organoid contours that were segmented. Scale bar = 50 µm.

**Supplementary Figure 2:**
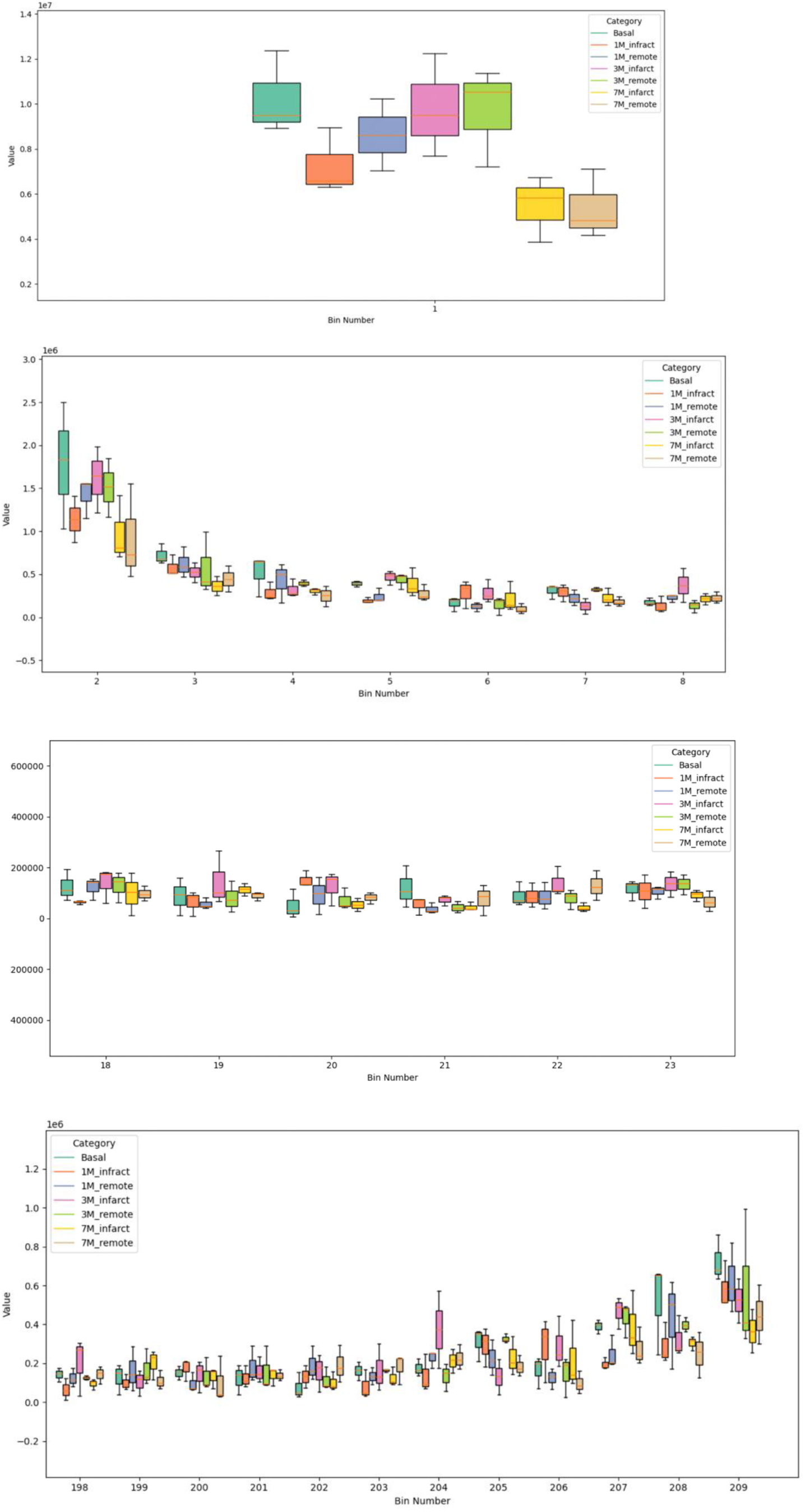
Box-whisker plots comparing the magnitudes of various frequency bins across the LCPC transform results of vascular networks. The LCPC transform of the vascular networks described in **Figure 9 and Figure 10** results in a set of frequency bins and their magnitudes that represent each image. Box-whisker plots of the magnitudes in each bin are used to compare all treatment groups, which identifies which bin best serves as an index for stratifying texture-density that was measured via the LCPC transform. Each box-whisker plot consists of n = 3 independent images.

**Supplementary Figure 3:**
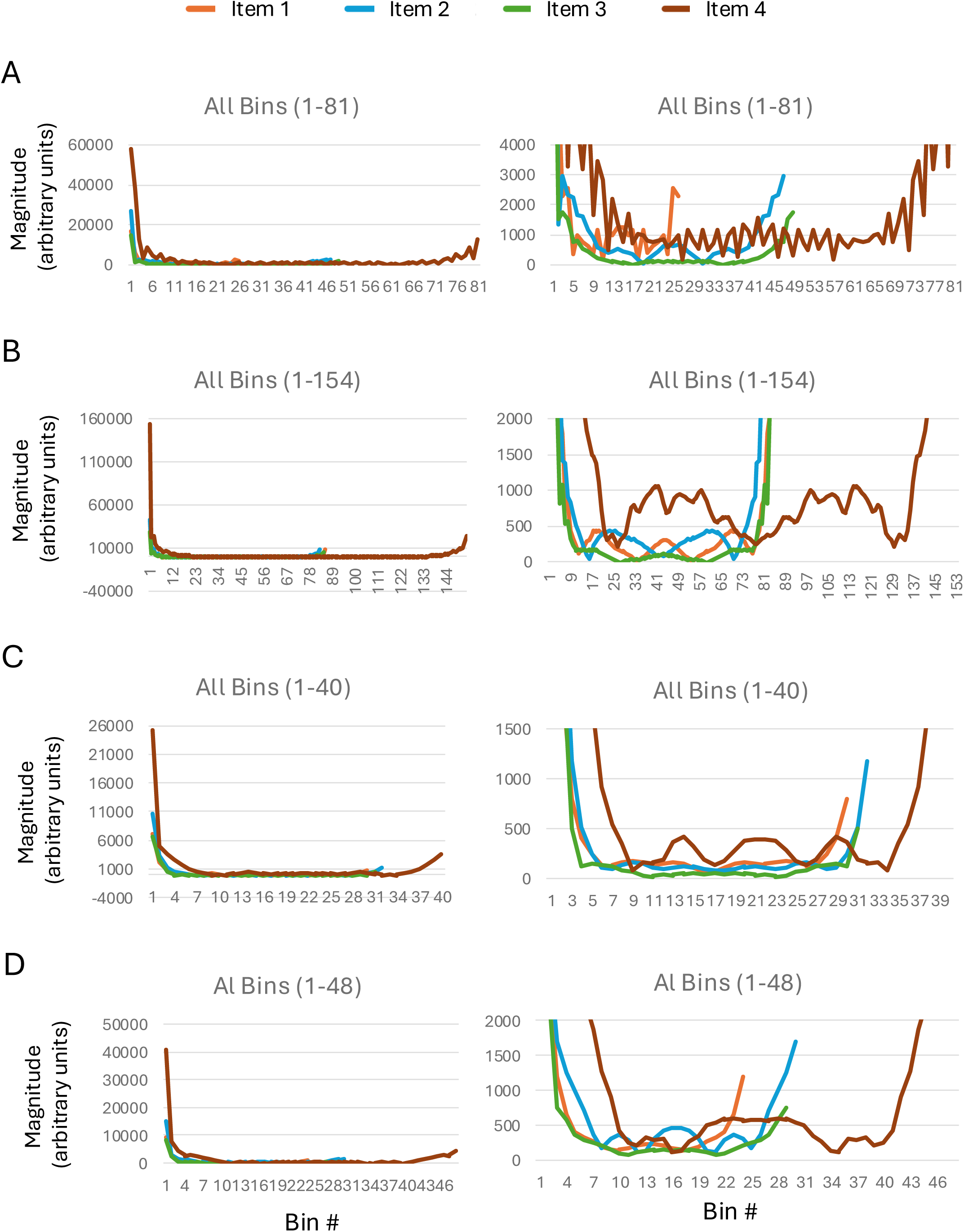
FFT spectra for shapes in Figure 6. The results of the FFT for the shapes in **Figure 6** are provided here so that readers can concretely understand the effect of altering the orientation of shapes on their spatial information. The left column of graphs has unaltered y-axes, while the right column has truncated axes such that the smaller bin values can be visualized. (**A**) Spectra from the parallel grid LCPC transform of the four shapes in **Figure 6C** as they are displayed, meaning no re-orienting. (**B**) Spectra from the parallel grid LCPC transform of the shapes in **Figure 6D**, which have been re-oriented as described in the text. (**C**) Spectra from the parallel grid LCPC transform of shapes in **Figure 6E**. (**D**) Spectra from the parallel grid LCPC transform of shapes in **Figure 6F** for comparing with those in **Figure 6E**.

**Supplementary File 1: Extensive details on best practices for the LCPC transform.** This supplementary file provides extensive discussion on best practices and further methodological notes. The outline of the document is as follows: **Section 1** - Spatial context that area, volume, perimeter, and roughness cannot capture. **Section 2** - Best practices for using the LCPC transform and interpreting results. **Section 3** - Understand nature’s inherent polarity to maximize the LCPC transform. **Section 4** - Interpreting FFT spectra resulting from the LCPC transform. **Section 5** - How to orient shapes before measuring them with the LCPC transform. **Section 6** - What is “Pure Shape” vs. “At Scale Shape”? **Section 7** - Methodological notes on statistical methods. **Section 8** - Methodological notes on cell culture and segmentation of organoids. **Section 9** - Parameters of the FFT for those who code their own LCPC transform.

