## Supplementary File 1 for "A Shape Analysis Algorithm Quantifies Spatial Morphology and Context of 2D to 3D Cell Culture for Novel Quantitation of Phenotypes"

**Supplemental File 1**

Nguyen et al. “A Shape Analysis Algorithm Quantifies Spatial Morphology and Context of 2D to 3D Cell Culture for Novel Quantitation of Phenotypes”. 2026

**Outline**

Section 1. Spatial context that area, volume, perimeter, and roughness cannot capture………Pg. 2

Section 2. Best practices for using the LCPC transform and interpreting results……………………Pg. 2

Section 3. Understand nature’s inherent polarity to maximize the LCPC transform……………..Pg. 3

Section 4. Interpreting FFT spectra resulting from the LCPC transform………………………………..Pg. 3

Section 5. How to orient shapes before measuring them with the LCPC transform……………Pg. 4

Section 6. What is “Pure Shape” vs. “At Scale Shape”?............................................................Pg. 5

Section 7. Methodological notes on statistical methods.…………………………………………………..Pg. 6

Section 8. Methodological notes on cell culture and segmentation of organoids…………………Pg. 6

Section 9. Parameters of the FFT for those who code their own LCPC transform…………………Pg. 7

**Section 1. Spatial context that area, volume, perimeter, and roughness cannot capture**
Biology and nature are full of spatial contexts that are crucial to proper function. Epithelial cells have inherent morphological spatial context, such as apical-basal polarity, to execute their function of forming barriers and directionally secreting products[36,37,38,39]. When cells migrate 2D cell culture, the leading edge of the cell has a distinct morphology compared to the trailing edge. This asymmetry in cell morphology during migration is essential for sensing the environment and for anchoring the cell for generating force in specific directions.

Another crucial spatial context in nature is the direction of gravity. Bone health, muscle health, and vascular health in astronauts become dysfunctional due to prolonged exposure to microgravity conditions[40,41]. Not surprisingly, cancer cells behave very differently in microgravity compared to being on Earth, including alterations in their morphology as single cells and in 3D colonies[42,43,44]. A third spatial context in cancer is that in situ pre-neoplasia and neoplasia can sprout from a base, or trunk, which is its central support structure. Pedunculated colon polyps are an example of this morphology. As the lesion regresses, it changes shape towards this central support structure, which is also its bridge to the stable vascular network that carries oxygen[45,46]. Traditional shape metrics used in cancer research and cell biology include measuring area, surface area, volume, perimeter, etc. While these metrics are informative, they cannot register spatial contexts that are obvious to humans. The apical-basal polarity of a cell in a lumen, the direction of gravity, and the direction of the central support structure is non-existent to traditional approaches that cannot register spatial context.

Traditional metrics also cannot distinguish gradations of shape change between common shape concepts, such as a square that is standing on one of its flat sides and a diamond that is the same square but rotated so that it stands vertically on one corner. Any intermediate forms resulting from rotating the square about its center to become the diamond are subjective interpretations. Are the intermediate steps “rotated squares” or “rotated diamonds”? Measuring area and perimeter to answer this question is pointless because the square and diamond are equivalent by these two metrics. Another shortcoming of traditional shape metrics is that they cannot distinguish between size and shape. A series of circles that gradually increase in size will exhibit increasing area and perimeter measures, even though the shape concept never changes, meaning the circles remain circles.

The existence of automated high-throughput cell culturing techniques that include imaging components means that morphology data is accessible as part of the research and development pipeline. However, without methods that robustly quantify subtle differences in morphology, it is difficult to discover unknown morphological subtypes and/or leverage the power of machine learning to correlate these subtypes with disease progression and treatment resistance.

**Section 2. Best practices for using the LCPC transform and interpreting results**
This manuscript describes only two grid systems for the LCPC transform, though many grid systems can be envisioned. It is important to note that while all grid systems can extract spatial information, some will provide cleaner data than others. Thus, users are encouraged to apply both the parallel and the radial grid systems to their data to determine optimal fit. For simplicity

and reproducibility, the open-source Python scripts only include a 360-degree radial grid system[17] and a parallel grid system[18], which will suffice for most applications. Again, all grid systems can extract spatial information, but some are better than others depending on the spatial contexts described in this manuscript. The aforementioned GitHub repositories contain links to YouTube videos that are tutorials on how to run the scripts on the associated practice images and how to do quality control checks on the outputs. It is highly recommended that users learn how and where to do quality control checks.

In general, the radial grid system is good for shapes that are radially symmetric, round, and closed (**Figure 1C**). However, it also works well for shapes that are open (meaning the contour has at least one disconnected region) but curved in on itself (like the letter "c"). This grid system also works for contours that consist of multiple layers or even contours that have multiple non-intersecting components. The parallel grid system, on the other hand, is good for shapes that are elongated and open (**Figure 2C**). Though it was previously stated that the radial grid system works well for closed shapes, closed shapes that are elongated sometimes yield cleaner results when assessed by the parallel grid system. Just as with the radial grid system, the parallel grid system can measure the spatial information of objects that consist of multiple non-intersecting components.

**Section 3. Understand nature's inherent polarity to maximize the LCPC transform**
Nature is inherently polar, which includes the shapes of cells and organoids. Cells can have apical-basal polarity and organoids can have asymmetries that indicate poles. Thus, orienting shapes in a systematic and objective manner before extracting spatial information via the LCPC transform will yield cleaner, more reproducible results. **Figure 6A** shows two shapes that are identical but mirror images of each other. Though they are the same shapes, because the parallel grid system always measures distances of intersections from the left side of the image, the parallel grid LCPC will yield different numbers for each image. The LCPC transform was designed to capture spatial context, so it is very sensitive to seemingly minor differences in spatial context. A better way to compare Shape 2 to Shape 1 in **Figure 6** is to flip Shape 2 horizontally, which changes the spatial orientation but doesn't change the shape, and then apply the parallel grid LCPC (as shown in **Figure 6B**). Compared to traditional shape metrics, such as area, volume, and perimeter, that are not affected by spatial orientation, the LCPC transform requires the user to think about how shapes should be oriented before measurement so that the optimal results are produced. The volume of liquid in an unopened can of soda does not change depending on whether the can is right-side up, upside down, or any other orientation. Volume is rotationally invariant, but the spatial context of nature, such as which side of an object is directly exposed to the sun, is often not.

**Section 4. Interpreting FFT spectra resulting from the LCPC transform**
The typical result of the LCPC transform, including the last FFT step, is a histogram of magnitudes as a function of position (**Figure 1A** Step 5). The first bin has the largest magnitude, and subsequent bins can exhibit various repeating patterns. **Figure 10** displays several hypothetical FFT results that have been observed. Each shape analyzed by the LCPC transform results in many frequency bins and their magnitudes that represent it, so using an unsupervised dimension

reduction technique, such as Principal Component Analysis (PCA), is useful for exploring which bins are responsible for stratifying the experiment groups most. However, a complementary approach is to derive a single-valued scalar index from the FFT results to represent each shape. These indices allow for the ease of employing descriptive statistics, such as the t-test or Wilcoxon Rank-Sum Test, to show differences in averages between experimental groups. They also allow for multiple distinct indices to be derived from the same FFT output, each of which *may* represent different morphological features in the shapes being analyzed. This ability to provide multiple indices that each represent a different feature of morphology is unprecedented compared to traditional metrics, such as measuring the area within an object, which provides no more than one value for area. **Figure 10A-10C** show patterns in bin magnitudes that can be used to create indices in the form of ratios. **Figure 10A** shows that there are three indices that can be derived based on the patterns highlighted by the dotted lines. Indices can be ratios between two bins, a sum of 2 or more bins, or a coefficient of a curve fitted onto multiple consecutive bins. **Figure 10D** shows hypothetical shape features that can correlate with different indices derived from the same FFT profile (data from actual experiments not shown). For an idea of how the FFT spectra changes due to shape orientation, **Supplementary Figure 3** shows actual FFT spectra of the shapes in **Figures 6C-6F**.

**Section 5. How to orient shapes before measuring them with the LCPC transform**

When measuring shapes with the LCPC transform, it is helpful to determine if the shapes should be oriented in a systematic way beforehand. **Figure 6C** shows four items that are shaped like the number "3" and arranged haphazardly. Unless there is a reason to keep each shape in its original orientation, the user should orient each shape like in **Figure 6D**. The rules for systematically aligning all items in **Figure 6C** into **Figure 6D** are: (1) The object is rotated such that the line that connects the opening of the "3" is vertical, (2) the wider part is on top, (3) and all objects have their concavity facing to the left so that the bulge of "3" pushes away from the left of the object where the parallel grid LCPC will place the reference baseline for measuring distance. Note that these rules require that Item 1 and Item 4 be both rotated and flipped horizontally, while Item 2 and Item 3 only need rotation. Flipping horizontally changes the spatial orientation of a shape, but not necessarily the shape. The user needs to determine what orientation rules to apply, and why, to every object, which will dramatically improve the cleanliness, and thus reproducibility, of the data.

It is important to note that the LCPC transform can be applied to sub-compartments of a shape instead of the whole shape. Morphologies in nature are highly complex, so scientists often assume insignificance for minor features of morphology to simplify the system being studied. However, the LCPC transform reveals that seemingly minor differences in morphology can be quantitatively measured and correlated with distinct biological function. Thus, **Figure 6E** shows how the wider bulge of the items in **Figure 6C** can be extracted and objectively compared. However, **Figure 6F** shows how the user can compare these wider bulges in the context of the alignment rules in **Figure 6D**. **Figure 6E** and **6F** are meant to highlight the fact that the user must determine what spatial context is relevant and why.

One of the limitations of traditional shape metrics, such as area and perimeter, is that they measure both the shape of an object and the size of an object at the same time. **Figure 7A** shows a series of check mark-shaped objects that increase in size but are the same shape concept. While the area and perimeter of each check mark change, the shape concept stays the same, but this is undetectable by area or perimeter. **Figure 7B** shows the concept of a square that becomes a diamond if it is rotated by 45 degrees. However, for any rotation between 0 and 45 degrees, the shape concept can be subjectively interpreted as either a "tilted square" or a "tilted diamond". Through this series of rotations, the area and perimeter of each object don't change while the shape concept does. The examples in **Figure 7A** and **7B** highlight the inability of traditional shape metrics (such as area, volume, perimeter, and surface area), and metrics that are rotationally invariant, in capturing notions that are obvious to humans.

**Section 6. What is "Pure Shape" vs. "At Scale Shape"?**

When measuring shapes, the LCPC transform can separate the effect of size from the effect of shape. In combination with an objective and systematic method of resizing objects, the LCPC transform can measure what will be called "pure shape", which is shape without the effect of size. **Figure 8** shows two objects that are different in both shape and size. If they are not resized before being measured by the LCPC transform, then they are being compared "at-scale", meaning at their original scales relative to each other. However, by applying an objective set of rules for rotation and resizing, the two objects can be transformed to a similar scale while keeping their original shape. **Figure 8** Step 2 shows how the longest internal line of each object is an objective morphological landmark by which the object can be rotated such that this line is horizontal. Step 3 of **Figure 8** then shows how the width of both objects can be resized to be 400-pixels wide while also constraining the aspect ratio so that the object is not skewed (not constraining the aspect ratio while resizing a square turns it into a rectangle). In this way, both objects now have the same width. 400-pixels wide is an arbitrary value written into the open-source script. Lastly, Step 4 of **Figure 8** shows an optional step of rotating the longest internal line within each object to be vertical, such that they get optimal coverage from the parallel grid LCPC transform. If the radial grid LCPC transform were applied, then this last rotation is unnecessary. However, it is suggested that the longest axes of objects be made vertical in case the user wants to also apply the parallel grid LCPC transform on the same objects for comparison with results from the radial grid LCPC transform. In the vertical position, the parallel grid LCPC transform applies the greatest coverage across the object because the parallel lines are stacked vertically.

Because traditional metrics, such as area, volume, and surface area, are agnostic to spatial context such as the direction of gravity, the direction of the strongest sunlight, and the axis of polarity, the LCPC transform provides a unique opportunity to insert spatial markers that capture these contexts. As described previously, how one rotates and orients a shape can also capture spatial context. **Figure 9A** shows another example of this in the form of cell migration in 2D culture. **Figure 9B**, however, is an example of how adding rationally placed information to the mask image allows the LCPC transform to capture spatial context in exquisite ways. **Figure 9B** shows a hypothetical tumor or pre-cancerous lesion that is anchored outside of its encasement. The lesion grows or regresses within the encasement, while the encasement does not change.

Thus, as the lesion grows away from, or regresses back to, its anchor due to therapeutic treatment, it does so relative to an encasement that remains static. This means that measures of area, perimeter, and volume (if the imaging context is 3D) would completely miss the fact that the spatial change in the lesion is happening within the context of an encasement. Why is this significant? In disease progression, the shape of cells, tissues, and organs do not change uniformly on all sides. They often grow away from or shrivel towards their central support structure. Furthermore, the "empty" space that they create relative to their encasement can have biological significance. Traditional shape metrics cannot capture the change of this "empty" space. **Figure 9C** shows two options for applying the parallel grid LCPC transform to the contour of the lesions. The orange line in **Figure 9B** represents the location of the encasement directly across from the leading edge of the lesion. Including this orange line in the mask of the lesion, as shown in **Figure 9C**, captures the fact that there is significantly more empty space between Specimen 1 and its encasement than for Specimen 2 and its encasement. The red dots on the green gridline in **Figure 9C** represent intersections between the gridline and the contour. Option 2 shows that the green line has four intersections instead of three in Option 1. This fourth intersection adds quantitative data the captures spatial context. The dotted lines in **Figure 9B** represent an objective and rational manner to include the spatial context of the encasement without adding too little or too much extra information.

**Section 7. Methodological notes on statistical tools employed and cell culture parameters**

When doing PCA on data from the LCPC transform, the data should first be median-centered instead of mean-centered, because the LCPC transform always results in a wide range of magnitudes and thus have what seem to be outliers. Bins 1, 2, and 3 are usually much greater in magnitude than the rest of the bins. However, the frequency domain of the FFT typically exhibits data like this, so labeling the extreme values as outliers is incorrect. Proper usage of PCA on data produced by the LCPC transform requires an awareness of the wide data scatter resulting from the FFT.

Cliff's Delta is used in place of a t-test or Wilcoxon Rank-Sum Test due to limited sample sizes and the non-parametric nature of the data. For n = 3 per group, there are 3 x 3 = 9 possible pairwise comparisons for Cliff's Delta: -1, -0.78, -0.56, -0.33, -0.11, 0.11, 0.33, 0.56, 0.78, 1. Delta > 0 indicates that the second set of values has shifted downward, while Delta < 0 indicates that second set has shifted upwards. Effect size labels: N = negligible ($|\delta| < 0.11$), S = small ($0.11 < |\delta| < 0.33$), M = medium ($0.33 < |\delta| < 0.56$), and L = large ($|\delta| > 0.56$)). In addition to the above statistical methods, the following methods were calculated using standard libraries in Python (v3.12.4): K-means Clustering (the value for K was determined using the elbow method), Silhouette Score, Chamfer Distance, Wasserstein Distance, and Average K-nearest Neighbor Distance (the value for K was determined as the square root of the sample size).

**Section 8. Methodological notes on cell culture and segmentation of organoids**

Organoids were generated from human breast epithelial tissue following a previously published protocol[19,20,21], under IRB-approved tissue collection protocols at University of California, San Francisco, and at Brigham and Women's Hospital. Surgical samples were processed on the day of surgery. In brief, tissues were digested for up to 2 hours with 1 µg / ml collagenase (Fisher, Cat. No. 17101015), then cell pellets were directly embedded in a 50 µl drop of basement membrane extract type 2 (Cultrex, Cat. No. 3532-005-02) in a dome in the center of a pre-warmed 24-well tissue culture plate (Greiner, Cat. No. M9312). This dome was then allowed to harden at 37 °C for 20 min and subsequently submerged in 500 µl of Type 2 Organoid Medium[19]. Plates were kept in humidified 37 °C / 5% CO2 incubators at ambient $O_2$ and organoid Medium was changed every 2-3 days. Organoids were passaged when confluent, depending on their growth rate, typically every 2-4 weeks, as previously described[19]. Images were obtained on an Echo Revolve Microscope, then important into ImageJ where individual structures were manually categorized and outlined depending on the number and density of cystic appearance of each organoid. Organoid boundaries were manually traced by an experienced investigator from the original microscopy images. See **Supplementary Figure 1** for examples of segmentations.

**Section 9. Parameters of the FFT for those who code their own LCPC transform**

It is important to note that the results of the LCPC transform cannot be reconstructed in reverse to derive the original shape of the input object. While the results of the FFT on typical discrete sinusoid data does allow for reconstruction, doing so on FFT data what was obtained on LCPC Transform results does not make sense. Reminder: the LCPC transform technically finishes before the FFT is applied as a last step, though the phrase “LCPC transform” in this manuscript is meant to include the FFT as part of the LCPC transform’s routine operation unless specifically stated otherwise. Reconstructing the discrete sinusoid wave that is the LCPC transform from the FFT spectra is doable, but because the LCPC transform sums all distances of intersections between the contour and a gridline for that gridline, the reconstructed wave won’t be the original shape.

FFTs were computed directly from the sampled data using NumPy's FFT implementation. No windowing, zero-padding, segment averaging, overlap, or multitaper spectral estimation procedures were applied. For FFT in the scripts provided, a sampling rate (a.k.a. sampling frequency) of $f_s$ = 40 Hz was chosen because the original way the sampling rate was picked was by dividing a 200-pixel-tall image by 5 pixels resulting in 40 horizontal lines. The FFT length (N) is the number of samples in the data; since no zero-padding is used, FFT length equals signal length. The frequency resolution (Δf) is $f_s$ / N = 40 Hz / N. For those who desire to rewrite the scripts to better fit their computational workflow, the scripts calculate a single-sided magnitude for the FFT. The GitHub repositories that contain the two LCPC scripts also contain practice input files and their associated output files. This is so that users and those who re-write the code have images that can be used to benchmarks to determine if their script produces the same results as the original scripts.

While the scripts for the LCPC transform are open-source and thus available for recreation and integration into existing computational pipelines, users who create their own version or modify the preset parameters should check to see if their results match those that are provided as part of this study. The GitHub repositories for the parallel grid and radial grid LCPC transform scripts contain several sets of practice images and their associated output files. The purpose of including both input and output files is so that users can check to make sure that their version of the script produces identical data as those provided in this study. Most users will adopt the preset parameters in the existing publicly available scripts, so those who alter or integrate the scripts for their own purposes should be aware that their data may no longer match everyone else's.
